# NINJA: an inducible genetic model for creating neoantigens *in vivo*

**DOI:** 10.1101/2020.01.09.900894

**Authors:** Martina Damo, Brittany Fitzgerald, Yisi Lu, Mursal Nader, Ivana William, Julie Cheung, Kelli A. Connolly, Gena G. Foster, Elliot Akama-Garren, Da-Yae Lee, Greg P. Chang, Vasilena Gocheva, Leah M. Schmidt, Alice Boileve, Josephine H. Wilson, Can Cui, Isabel Monroy, Prashanth Gokare, Peter Cabeceiras, Tyler Jacks, Nikhil S. Joshi

**Author notes:** Authors contributed equally to this work (Martina Damo, Brittany Fitzgerald). Authors contributed equally to this work (Tyler Jacks, Nikhil S. Joshi).

## Abstract

Mouse models with inducible neoantigens have been historically difficult to generate because of leaky expression of antigens in the thymus, which causes central tolerance in developing CD8 and CD4 T cells. Attempts to resolve this problem using existing genetic tools have been unsuccessful. We developed the iNversion INducible Joined neoAntigen (NINJA) mouse model that uses RNA splicing, DNA recombination, and three levels of regulation to prevent neoantigen leakiness and allow tight control over the induction of neoantigen expression. We describe the development of these genetic tools and their use for obtaining tumor cell lines with inducible neoantigen expression. Moreover, we show that the genetic regulation in NINJA mice bypasses central and peripheral tolerance mechanisms and allows for robust endogenous CD8 and CD4 T cells responses upon neoantigen induction in peripheral tissues. Thus, NINJA fills a long-standing gap in the field and will enable studies of how T cells respond to defined neoantigens in the context of peripheral tolerance, autoimmune diseases, and developing tumors.

---

The field of immunobiology has learned a great amount about how T cells function in the context of infections, cancer, and autoimmune responses using animal models. Yet, one area that has been difficult to study has been the modulation of endogenous T cells in peripheral tissues. One reason for this is that mice have a robust system of central tolerance established in the thymus that results in the “pruning” of the naïve T cell repertoire by eliminating or tolerizing CD8 and CD4 T cells with T cell receptors (TCRs) that recognize self-antigens^1–5^. These processes are crucial for maintaining tolerance but present a major engineering challenge for investigators interested in studying peripheral tolerance or in expressing inducible neoantigens from genome-encoded loci.

There have been several previous attempts to develop animal models that allow one to induce neoantigens in peripheral cells in a manner that bypasses central tolerance. These include efforts to suppress the transcription/translation of a downstream neoantigen using genetic elements such as lox-STOP-lox (LSL) elements or tet-inducible promoters ^6–11^. In the OFF state (pre-induction), these greatly reduce neoantigen expression, often to levels that are below the limit of detection by conventional methods. Yet, these methods often do not abrogate protein production completely, and there is a residual low level of “leaky” expression that, in the thymus or periphery, is sufficient to generate central or peripheral tolerance ^7^. As a result, it has been difficult to generate mice with inducible genes that are lethal (*i.e.,* LSL or tet-inducible Diphtheria toxin), as these mice can have phenotypes from their leaky expression in sensitive tissues ^12, 13^.

Central tolerance generally manifests as an incomplete or absent antigen-specific endogenous T cell response after infection with an antigen-expressing virus/bacteria or challenge with an antigen-expressing tumor cell line ^7^. Alternatively, central tolerance occurs after crossing neoantigen-inducible mice to transgenic (Tg) mice that express TCRs specific for the neoantigen ^14, 15^. Here, central tolerance can take on many forms including outright deletion of neoantigen-specific T cells, marked-downregulation of TCR avidity, and the increased development of natural regulatory T cells (Tregs) ^7–11, 16^. Transfer of TCR Tg CD8 or CD4 T cells into inducible neoantigen mice has also been used to bypass central tolerance, but this can result in the activation and even tolerance of the transferred cells prior to antigen induction ^7, 11^. This has required groups to provide inflammatory signals or infections coupled to antigen delivery to break antigen-specific tolerance, thus resulting in T cells attacking antigen-expressing self-tissues ^17–,19^, which can perturb the goal of studying how natural tolerance develops or is broken in peripheral tissues.

There is a clear need in the field to develop technology that allows for tight control over inducible expression of the neoantigens from germline encoded loci. To this end, we engineered the iNversion INducible Joined neoAntigen (NINJA) mouse to be compatible with Cre-inducible genetically-engineered mouse models of cancer like the “KP” model (Kras LSL-G12D;p53 flox/flox, ^20^). The NINJA allele consists of two modules. The first one is a “neoantigen” module that uses DNA inversion to permanently induce neoantigen expression. DNA inversion is regulated by Flippase (FLPo) ^21, 22^, which is encoded by a second “regulatory” module. In the regulatory module, FLPo is regulated at three levels requiring Cre-recombinase mediated recombination to “poise” the module, followed by doxycycline (D) and tamoxifen (T) to induce FLPo expression and activity, respectively.

## Results

### Design of the spliced neoantigen module

To generate a module encoding a neoantigen that would not have leaky expression in the OFF state, we reasoned that the DNA sequences encoding the full-length neoantigen should not exist within the genome prior to induction. Thus, we conceived of a gene in which the neoantigen-encoding DNA sequences were split across three exons, and the second exon was inverted relative to the first so that the gene could not transcribe/translate the full-length peptide antigen in the OFF state (**Fig 1A**). Induction would be triggered by inversion of the second exon, which we determined would be achieved using FLPo and “non-compatible” FRT sites ^21, 22^.

**Figure 1.**
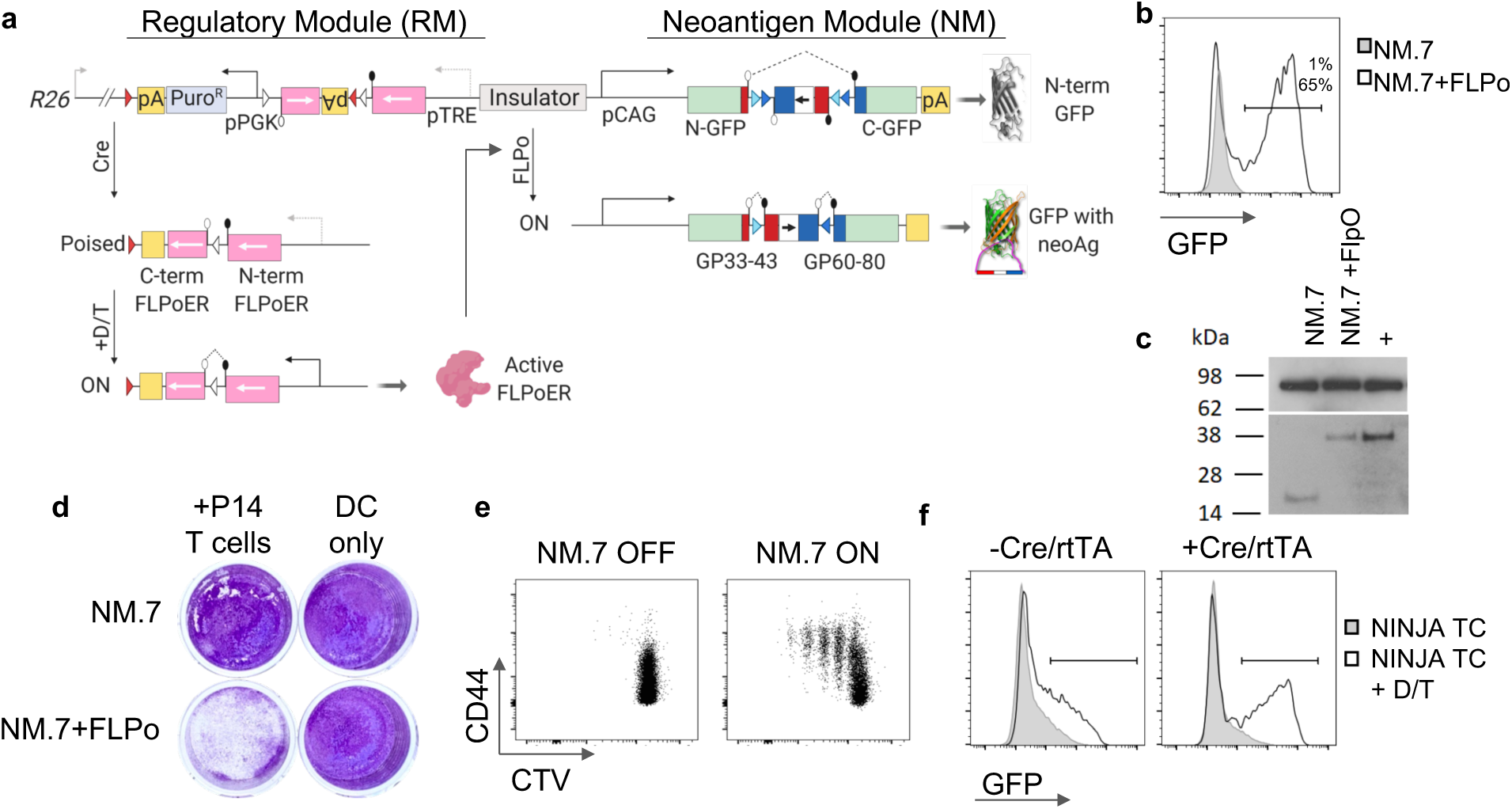
Design and development of NINJA targeting construct. **a,** Schematic of NINJA targeting construct (NINJA TC) after integration into Rosa26 (R26) locus. NINJA contains regulatory (RM) and neoantigen (NM) modules. Schematic shows modules after Cre or FLPo recombinase and doxycycline/tamoxifen (D/T) exposure. (Red and white fill arrow pairs are non-compatible loxP sites. Black and white ball and stick lines are splice sites, and black or grey arrows indicate promoters) **b-c,** FLPo exposure is required for GFP expression from NM. 293T cells were transfected with the indicated constructs and assessed by FACS or western blotting. **b,** Histogram shows GFP expression 72 hours after transfection with NM.7 alone (gray, filled) or NM.7 and FLPo (line). Percent GFP+ for each is indicated. **c,** Western blotting for GRP94 (control, top panel) or N-terminal GFP (bottom panel) on lysates from 293T cells. Positive control (+) is cell lysate from KP-C4A3D6 after FLPo (see Fig. 2). **d-e,** NM is only immunogenic after FLPo exposure. DC2.4 cells were transfected with NM.7 or NM.7+FLPo and cultured with or without naive Cell Trace Violet (CTV)-stained CD8 P14 T cells. **d,** Tumor cell wells were stained with crystal violet four days later. **e,** FACS plots show CTV dilution and CD44 expression on CD8+ Va2+ T cells. **f,** NINJA TC expresses GFP after Cre, rtTA, and D/T treatment. 293T cells were transfected with NINJA TC with and without plasmids expressing Cre and rtTA. Cells were then cultured with or without D/T. Histograms show GFP expression from the 293T cells with the indicated conditions.

First, we identified neoantigens that had immunological tools for their analysis (*i.e.,* MHC class I and class II tetramers, TCR Tg mice, *etc.*) **and** had sequences that could be engineered to contain in-frame splice donor and acceptor sites (**Fig S1A**). Thus, we settled on well characterized antigens from the glycoprotein (GP) of the lymphocytic choriomeningitis virus (LCMV), which is recognized by H2-D^b^-restricted CD8 T cells (GP33-41: **KA**VYNFATCGI) and H2-I-A^b^-restricted CD4 T cells (GP67-77: IY**KG**VYQFKSV) [splice sites were inserted between the amino acids noted in bold]. GP33-41 and GP66-77 are recognized by P14 and SMARTA CD8 and CD4 TCR Tg mice, respectively. *In silico*, we inserted RNA splice sites into the DNA and created two introns (**Fig S1B**). We also inverted the sequences encoding exon 2, so that in the OFF state, the construct would splice directly from exons 1 to 3. To allow for induction of neoantigens in response to the recombinase FLPo, we inserted non-compatible FRT sites (F3 and FRT Wt; ^22^) to flank the inverted exon 2 sequence so that exposure to active FLPo would cause a permanent inversion of exon 2 (**Fig S1C**, ^23^).

Over the course of 7 iterative versions of the neoantigen module (NM.1-NM.7) (**Fig S2A**), we refined the design to ensure that the constructs had high fidelity before and after FLPo-mediated recombination at the DNA, RNA and protein levels (**Fig S2, S3,** and data not shown). We also modified the construct so that inducible neoantigen was incorporated into the coding sequence of GFP in such a way that the non-fluorescent N-terminal portion of GFP was produced when the neoantigen module was OFF (before recombination) and full-length fluorescent GFP containing the neoantigens was produced when the module was ON (after recombination; **Figs 1A** and **S3**). Finally, we confirmed that the construct was unable to produce a cross-tolerizing GP33-43 peptide when the neoantigen module was in the OFF state (**Fig S4**).

### Validating the final neoantigen module construct

To test the final neoantigen module construct (NM.7) for functionality, we transfected 293T cells with the construct in the presence or absence of a plasmid encoding FLPo. After 72 hours, flow cytometric (FACS) analysis of the transfected cells showed robust GFP fluorescence in 293T cells transfected with the neoantigen module and FLPo, but no detectable fluorescence when FLPo was not co-transfected (**Fig 1B**). This corresponded with a change in the size of the GFP protein from 27 kDa to 32 kDa (**Fig S2C**). We also isolated the DNA from 293T cells after co-transfection with FLPo and confirmed that FLPo had faithfully recombined the neoantigen module by PCR and DNA sequencing (data not shown).

Next, we assessed the ability of the constructs to elicit responses by GP33-specific CD8 T cells. For this, we transiently transfected a dendritic cell line (DC2.4) with either GFP (negative control),the NM.7 construct or the NM.7 construct + FlpO. We cultured the transfected DC2.4 cells for 4 days alone or with cell-trace violet (CTV)-labeled CD8 T cells from GP33-43-specific P14 TCR Tg. In this assay, if P14 CD8 T cells recognize GP33-43 presented by DC2.4 cells, they become activated (dilute CTV and increase CD44 expression) and kill the adherent DC2.4 cells (leading to loss of crystal violet staining). In the GFP control (not shown) and NM.7 transduced wells, the P14 T cells remained naïve and the DC2.4 cells formed a confluent monolayer (**Fig 1D**). By contrast, when DC2.4 cells were transfected with NM.7+FLPo, P14 T cells were activated and proliferated (**Fig 1E**) and killed the DC2.4 cells (**Fig 1D**). Together, these data demonstrate that the NM.7 neoantigen module is non-immunogenic in the OFF state and immunogenic in the ON state after FLPo recombination.

### Design of the regulatory module

We designed the regulatory module to contain a Cre-inducible FLPo that is fused to amino acids 251-596 of the human estrogen receptor (ER) (FLPoER) and is expressed under the doxycycline-inducible pTRE-tight promoter (**Fig 1A** **and S5**). Similar to the neoantigen module, in the regulatory module the DNA encoding the FLPoER sequence has been spliced and split into two exons in opposite orientations and requires Cre-recombinase-mediated inversion to become able to produce a functional protein (**Fig 1A**). After Cre, the regulatory module is placed in the “poised” state, and subsequent FLPo activity requires D/T treatment. For this to happen, cells must also express the reverse tetracycline transactivator (rtTA), adding an extra potential layer of specificity (*i.e.,* tissue-specific promoters expressing rtTA).

### Assembly and testing of NINJA targeting construct

We inserted the regulatory and neoantigen modules into a *Rosa26* targeting construct such that they were oriented in opposite directions (**Fig 1A** and **S6**). This was done to avoid transcriptional interference in the regulatory module from the endogenous *Rosa26* locus promoter. To minimize the impact of the CAG promoter (in the neoantigen module) on the pTRE promoter in the regulatory module, we inserted the 2x chicken gamma globulin (CGG) insulator between the regulatory and neoantigen modules. This construct was assembled in a *Rosa26* targeting vector and will henceforth be referred to as the NINJA targeting construct (NINJA TC).

We next tested the NINJA TC in 293T cells. In the absence of Cre the neoantigen module remained OFF (GFP-), even when the cells were treated with D/T. By contrast, when cells were co-transfected with the full construct and Cre and rtTA, it was possible to turn on the neoantigen module (GFP+) with D/T treatment (**Fig 1F**). These data confirm that the targeting construct functions as designed and induces neoantigens after Cre, doxycycline, and tamoxifen treatment.

### Generating a NINJA-containing lung adenocarcinoma cell line

To test the functionality and fidelity of the components of the regulatory and neoantigen modules in the context of the *Rosa26* locus, we used a murine KP lung cancer cell line to generate a knock-in cell line containing the NINJA targeting construct (KP-C4) (**Fig 2A**). Proper targeting of the NINJA construct to the Rosa26 locus was verified by PCR (data not shown). We then transduced our KP-C4 cells with an rtTA-encoding retrovirus (KP-C4A3) and infected them with a recombinant Cre-expressing adenoviral vector (Ad-CRE) to cause a recombination in the regulatory module and place it *permanently* in the poised state (KP-C4A3D6) (**Fig 2A**). We verified that the neoantigen module in the KP-C4A3D6 cell line remained OFF for over 20 passages in culture (based on GFP fluorescence, not shown). However, when the KP-C4A3D6 cell line was infected with a recombinant adenoviral vector expressing FLPo (Ad-FLPo, positive control) or treated with doxycycline and 4-hydroxytamoxifen (4-OHT), these cells turned ON the neoantigen module and expressed GFP (**Fig 2B**).

**Figure 2.**
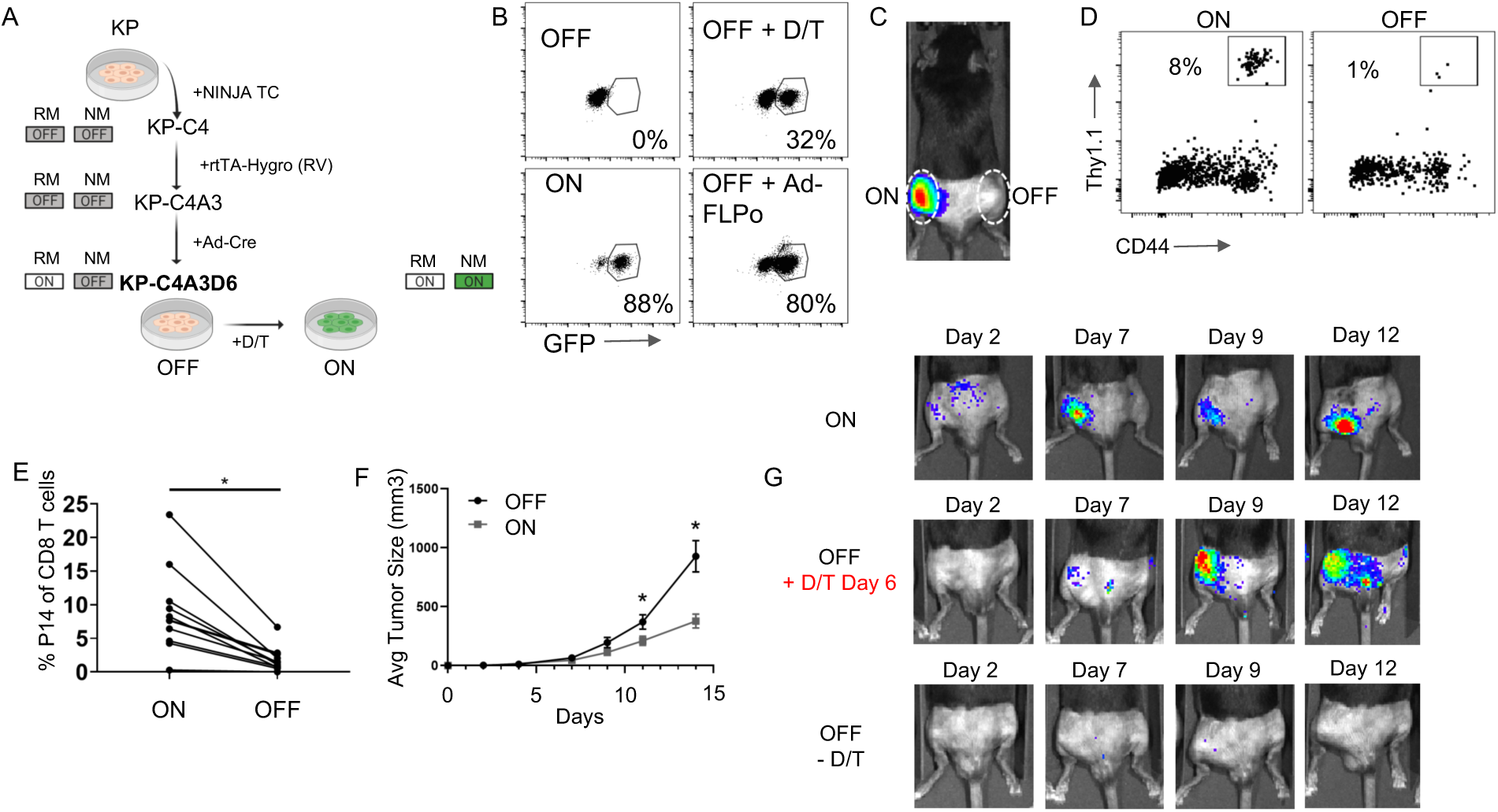
Stable NINJA cell line generates antigen specific T cell response following neoantigen induction. **a,** Schematic depicting generation of the “poised” KP-C4A3D6 cell line, which turns on neoantigen in response to D/T. b. FACS plots show GFP expression by untreated KP-C4A3D6 cells (OFF), or KP-C4A3D6 cells 72 hours after D/T treatment or infection with Ad-FLPo. Plots also show FACS-sorted GFP+ KP-C4A3D6 cells (ON). **c-f,** KP-C4A3D6 cells only elicit tumor-specific CD8 T cell responses *in vivo* when ON. KP-C4A3D6 OFF and ON cells were intramuscularly transplanted bilaterally into in C57Bl/6 recipients (25k-200k cells, total n = 24, 3 experimental repeats). Mice also received naive luciferase+ Thy1.1+ P14 CD8 T cells. **c,** Representative IVIS image shows signal from P14 CD8 T cells in ON and OFF tumors in the same mouse on days 9-11 after transplant. **d,** FACS plots show presence of intratumoral Thy1.1+ CD8+ P14 CD8 T cells and expression of CD44 from ON and OFF tumor from representative mouse. **e,** Graph shows the frequency of intratumoral Thy1.1+ CD8+ P14 CD8 T cells from paired ON and OFF tumors (connected by line). *, p<0.0006 (paired T test, n = 11). **f,** Graph shows average tumor measurements of tumors in ON and OFF legs +/-SEM (n = 14). **g,** In vivo activation of neoantigens triggers tumor-specific CD8 T cell response. KP-C4A3D6 OFF and ON cells were intramuscularly transplanted into C57Bl/6 recipients that contained naive luciferase+ Thy1.1+ P14 CD8 T cells. IVIS images show signal from P14+ CD8 T cells on indicated days. Mice in middle panels were treated with D/T starting on day 6 post tumor inoculation. (n > 12, 2 experimental repeats, n = 10/12 of treatment group responding).

To test immunogenicity, we sorted a pure GFP+ population of neoantigen expressing (ON) KP-C4A3D6 cells and compared these cells *in vivo* with KP-C4A3D6 that had never been exposed to D/T (OFF, **Fig 2B**). We implanted the ON and OFF cells intramuscularly (I.M.) in opposite legs of C57Bl/6 (B6) mice along with adoptively transferred firefly-luciferase (fLuc)-expressing P14 CD8 T cells. This allowed us to track the accumulation of neoantigen-specific CD8 T cells in the tumors by *in vivo* bioluminescence imaging (IVIS). After 11 days, we observed accumulation of fLuc+ P14 CD8 T cells in the neoantigen-expressing ON tumors, but not in the neoantigen-negative OFF tumors (**Fig 2C**). Infiltration of neoantigen-specific P14 CD8 T cells into neoantigen-expressing tumors was confirmed by FACS (**Fig 2D**) and we also noted that the infiltrating T cells had a PD-1^hi^ phenotype, consistent with local cognate antigen exposure (**Fig 2E** and data not shown).

One of the unique advantages of the NINJA system is the ability to induce the expression of neoantigens in palpable tumors with *in vivo* D/T treatment. To test this, P14 chimeric mice were implanted with KP-C4A3D6 cells (I.M.) and were subsequently treated with D/T beginning 6 days post-transplant. T cell responses were only observed in D/T treated mice or ON controls, with responses visible shortly after D/T treatment (**Fig 2G**). Together, these data confirmed that NINJA is a system for tight regulation and induction of neoantigen expression.

### Baseline “leaky” neoantigen expression not detected in NINJA mice

Having successfully generated and tested the NINJA targeting construct, we next generated Rosa26 targeted NINJA mice on a B6 genetic background (**Fig. S7**). NINJA mice were born at the expected mendelian frequencies and had no distinguishing phenotypes compared to wild-type B6 mice. We first assessed whether there was baseline leaky expression of neoantigens in NINJA mice by looking at the frequency of GFP+ cells in their peripheral blood mononuclear cells (PBMCs; **Fig 3A**). As positive controls, we bred NINJA mice to FLPo Tg mice to generate lines that had germline recombination of the neoantigen module and therefore ubiquitous constitutive expression of the NINJA neoantigens (NINJA-F). We also generated NINJA x Cre Tg (NINJA-C) mice, where the regulatory module of NINJA is maintained in the poised state in all cells throughout the body. In NINJA-F mice, all PBMCs were GFP+, while no PBMCs were GFP+ in NINJA mice, suggesting there is no leaky neoantigen expression in NINJA. Interestingly, in NINJA-C mice, >99% of PBLs remained GFP-, although we were able to detect a very small fraction of GFP+ cells (<1%) (**Fig 3A**; note, this fraction increases with age). These data suggest that the NINJA allele in mice remains very tightly regulated, without detectable neoantigen expression, except under extreme circumstances where regulatory elements are compromised.

**Figure 3.**
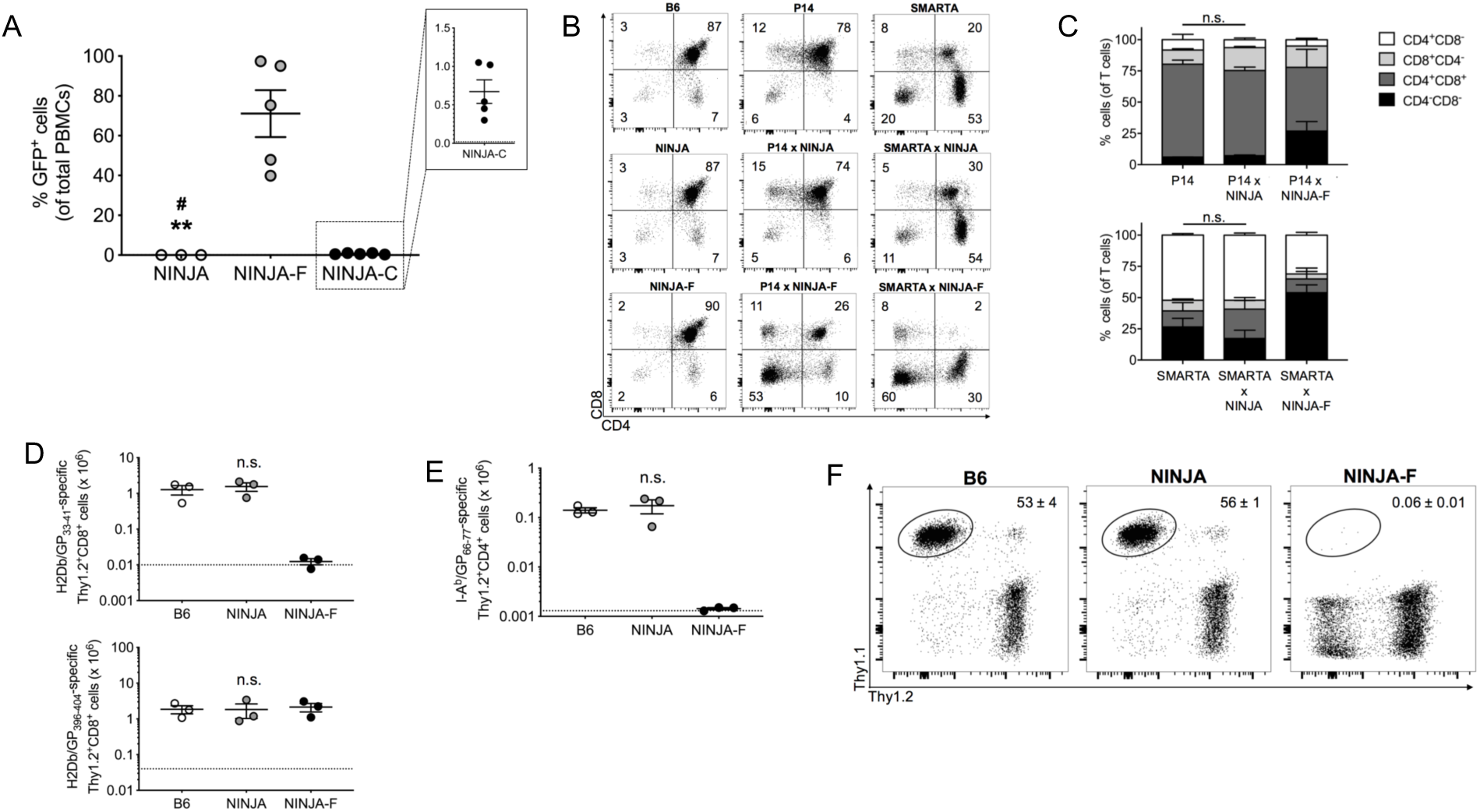
Central tolerance is bypassed in NINJA mice. **a,** No neoantigen expression at baseline in NINJA mice. Graph shows the frequency of NINJA-expressing GFP^+^ peripheral blood mononuclear cells (PBMC) isolated from NINJA, NINJA-F and NINJA-C mice (determined by flow cytometry). Dotted line represents background fluorescence signal from wild-type B6 PBMCs. Representative experiment (*n* = 15). **\*\*** *P* < 0.01 by unpaired *t* test for comparison of NINJA vs. NINJA-F; **#** *P* < 0.05 by unpaired *t* test for comparisons of NINJA vs. NINJA-C. **b-c,** NINJA mice bypass central tolerance towards neoantigens. **b,** FACS plots show CD4 and CD8 staining of Thy1.2+ thymocytes of the indicated mouse strains. Representative mice are shown (*n* = 10). Numbers represent frequency of gated cell populations. **c,** Graph shows quantification of data from **b**. **d-e,** NINJA mice have normal CD8 and CD4 T cell responses to LCMV infection. Graphs showing the number of **d,** H2Db/GP_33-41_-specific (top) and H2Db/GP_396-404_-specific (bottom) Thy1.2^+^CD8^+^ cells or **e,** I-A^b^/GP_66-77_-specific Thy1.2^+^CD4^+^ cells from the spleen on day 8 after LCMV-Armstrong infection. Representative experiment (*n* = 9). Dotted lines represent background tetramer staining as determined on splenocytes from untreated B6. Graphs in **a-e** show average values ± SEM. n.s., not significant by unpaired *t* test. **f,** T cells are not tolerized by environment of NINJA mice. Thy1.1^+^/1.1^+^ P14 cells were adoptively transferred into B6, NINJA and NINJA-F mice. 2 weeks later, mice were infected with LCMV-Armstrong. FACS plots show the frequency of P14 T cells on day 8 post infection from spleen.

### NINJA mice are not tolerant to their neoantigens

To test NINJA mice for central tolerance towards GP33-41 and GP66-77, we bred them to P14 CD8 and SMARTA CD4 TCR Tg mice respectively, and analyzed developing thymocytes. Similar to parental P14 or SMARTA mice (negative control), P14 and SMARTA x NINJA mice had normal development of CD8 and CD4 T cells, while severe defects in the development of CD8 and CD4 T cells became evident in P14 and SMARTA x NINJA-F mice (**Fig 3B-C**).

Consistent with their lack of central tolerance, NINJA mice had robust GP33-41-specific CD8 and GP66-77-specific CD4 T cell responses to acute LCMV infection, which were indistinguishable from the responses developed by control B6 mice, while NINJA-F mice were completely tolerant to the cross-reactive antigens (**Fig 3D-E**). Finally, to confirm that the peripheral environment of NINJA mice did not induce tolerance, we “parked” adoptively transferred 10^6^ P14 CD8 T cells in B6, NINJA or NINJA-F mice and infected them with acute LCMV after 14 days. The transferred P14 CD8 T cells were tolerized in the environment of the NINJA-F mice, but instead developed robust responses in both B6 and NINJA mice (**Fig 3F**). Together, these data demonstrate that in the absence of neoantigen induction, neoantigen-specific CD8 and CD4 T cells in NINJA mice are not being centrally or peripherally tolerized, as was seen in previous models.

### Neoantigen module is functional in cells from NINJA mice

To test the functionality of the neoantigen module in NINJA mice, we generated bone marrow derived dendritic cells (BMDCs) from NINJA mice and infected them with either a recombinant GFP-expressing adenoviral vector (Ad-GFP, control) or Ad-FLPo. We then used Ad-GFP-treated or Ad-FLPo-treated NINJA BMDCs to immunize B6 recipients that were previously adoptively transferred with 25,000 Thy1.1/1.1+ P14 CD8 T cells and Thy1.1/1.2+ SMARTA cells. The P14 CD8 T cells and the SMARTA CD4 T cells responded to the immunization by expanding and becoming detectable by FACS only in the mice that received Ad-FLPo-treated neoantigen-expressing NINJA BMDCs (**Fig 4A**).

**Figure 4.**
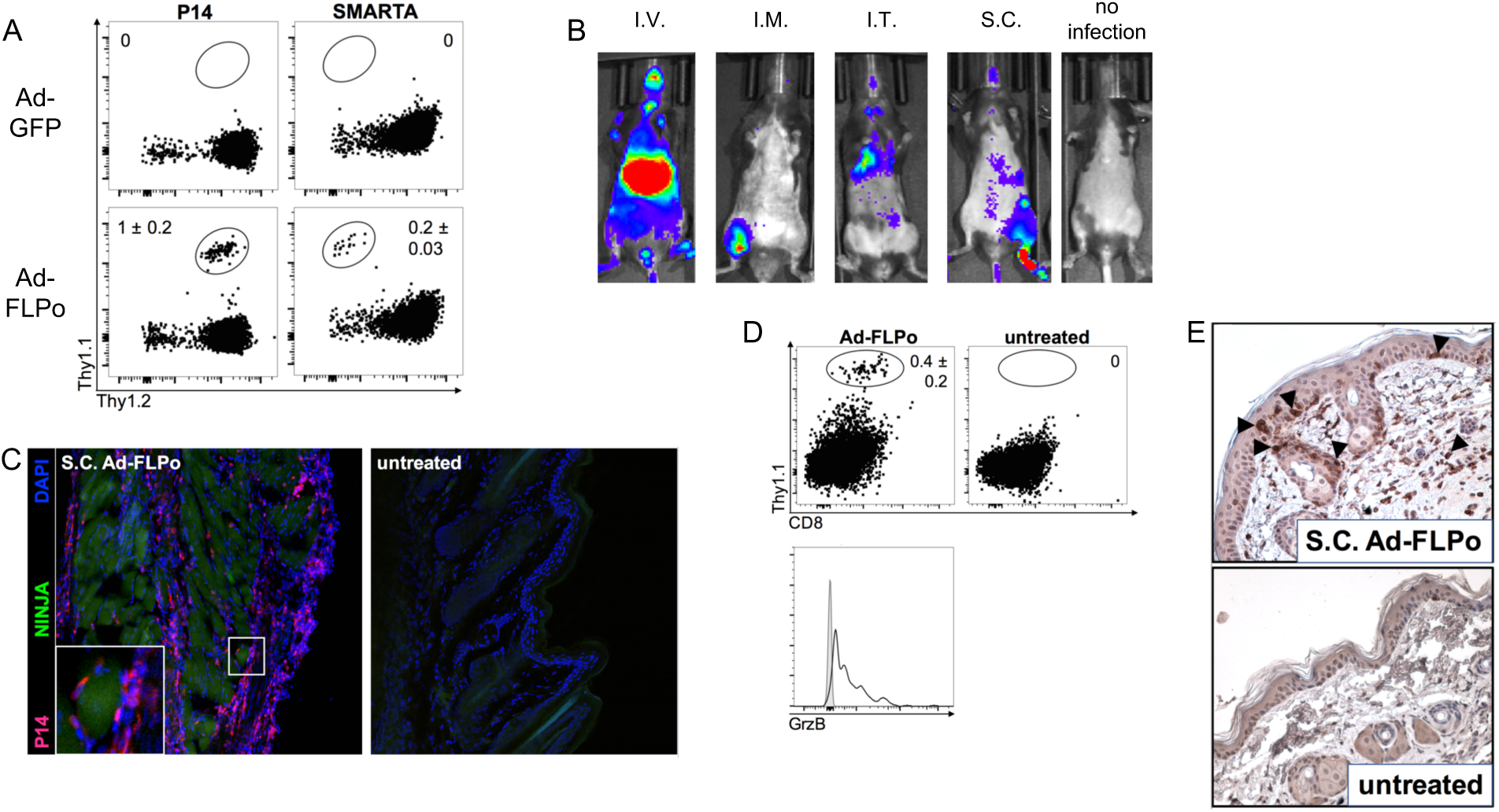
In vivo activation of P14 and SMARTA cells in NINJA mice after local infection with Ad-FLPo. **a.** Activation of neoantigen expression in NINJA BMDCs stimulates neoantigen-specific CD8 and CD4 T cell responses. BMDCs were generated from NINJA mice and infected with Ad-GFP (control) or Ad-FLPo to turn on NM. BMDCs were used to immunize B6 mice (S.C.) that contained adoptively transferred thy1.1^+^/1.2^+^ P14 CD8 T cells and thy1.1^+^/1.1^+^ SMARTA CD4 T cells. Dot plots show frequency of P14 (left) or SMARTA (right) T cells from the indicated mice from a representative experiment (*n* = 11). **b**. Expansion of neoantigen-specific CD8 T cells is controlled by site of antigen induction. fLuc^+^ Thy1.1+ P14 CD8 T cells were adoptively transferred into NINJA mice that were subsequently infected I.V., I.M., I.T. or S.C. (footpad) with Ad-FLPo. IVIS images show the locations of P14 T cells 8 days after infection. Representative mice are shown (*n* = 3). **c**. Neoantigen-specific T cells interact with neoantigen expressing skin cells after Ad-FLPo infection. dsRED^+^ P14 CD8 T cells were adoptively transferred into NINJA mice that were subsequently infected with Ad-FLPo S.C. in the footpad (left) or left untreated (right). Confocal microscope images show GFP^+^ NINJA-expressing cells (green), P14 T cells (red) and DAPI (blue) 8 days after infection. Representative images (*n* = 3). **d-e**, Neoantigen-specific CD8 T cells acquire effector functions after antigen induction in footpad. **d**, Histogram shows granzyme B expression by Thy1.1^+^ P14 CD8 T cells from Ad-FLPo (top left) infected or untreated (top right) NINJA mice on day X after infection Representative mice from X experiments (*n* = 6). **e**, Immunohistological sections of the footpad stained with anti-CD3 (brown). Images show epidermal/dermal thickening and infiltration by CD3^+^ cells (arrows) Representative images (*n* = 3). Average values ± SEM are shown in **a** and **d**.

### Neoantigen-specific T cells respond to *in vivo* neoantigen induction in NINJA

We next tested whether fLuc+ P14 CD8 T cells would respond to *in vivo* induction of neoantigens in NINJA, after infection with 10^7^ PFU Ad-FLPo in the footpad (subcutaneous, S.C.), muscle (intramuscular, I.M.), liver (intravenous, I.V.), or lungs (intratracheal, I.T.). In all cases, by day 8 post-infection we could observe by IVIS accumulation of fLuc+ P14 T cells specifically at the infected site (**Fig 4B**). Moreover, this system allowed us to track the kinetics and location of T cell expansion after self-neoantigen induction, which demonstrated that T cells were first activated in the tissue-draining lymph nodes (S.C., I.M., I.T.) or spleen (I.V.) and then migrated to the sites of neoantigen expression (**Fig S8** and data not shown). This suggested that in NINJA mice, soon after infection with Ad-FLPo, neoantigens are being expressed in the treated tissue and functionally presented in local draining organs, leading to activation of neoantigen-specific T cells.

To determine if the neoantigen-specific T cells were interacting with neoantigen-expressing cells in the tissues, we transferred dsRed-expressing P14 CD8 T cells into NINJA mice and 24 hr later infected them in the footpad with 10^7^ PFU Ad-FLPo. 8 days after infection, we observed dsRed+ P14 T cells localized in the infected footpad in direct proximity to GFP+ skin cells (**Fig 4C**). Moreover, P14 T cell accumulation in the infected footpad and acquisition of T cell lytic effector functions (Granzyme B expression) were confirmed by FACS (**Fig 4D**). Finally, signs of immunopathology were detected in the infected tissue, as shown by the infiltration of CD3+ cells and thickening of epidermal and dermal layer of the skin as compared to non-infected footpad (**Fig 4E**). Together, these data show that *de novo* induction of self-neoantigens in the context of an infection with a non-replicating adenoviral vector can trigger a potent effector T cell response in NINJA mice.

### Endogenous effector T cell responses against self-neoantigens

To determine whether *in vivo* induction of neoantigens could trigger endogenous neoantigen-specific CD8 and CD4 T cell responses, we infected NINJA mice in the footpad with 10^7^ PFU Ad-FLPo and analyzed mice 8 days later. We observed robust self-neoantigen-specific responses from both endogenous GP33-41-specific CD8 (**Fig 5A**) and GP66-77-specific CD4 (not shown) T cells. Moreover, the T cells were polyfunctional, as indicated by their expression of Granzyme B and pro-inflammatory cytokines IFNγ and TNFα (**Fig 5B**). Signs of a robust anti-tissue response with consequent immunopathology localized at the infected footpad were also detected (**Fig 5C**). Note, we performed similar experiments in NINJA mice after I.V., I.M., or I.T. administration of Ad-FLPo and in all cases tissue-restricted immune responses could be detected (not shown). These data confirm that *in vivo* activation of the NINJA neoantigen module in the context of adenoviral vector infection results in strong endogenous self-neoantigen-specific T cell responses, in line with the fact that neoantigen-specific T cells in NINJA mice are truly naïve for their cognate antigens prior to induction.

**Figure 5.**
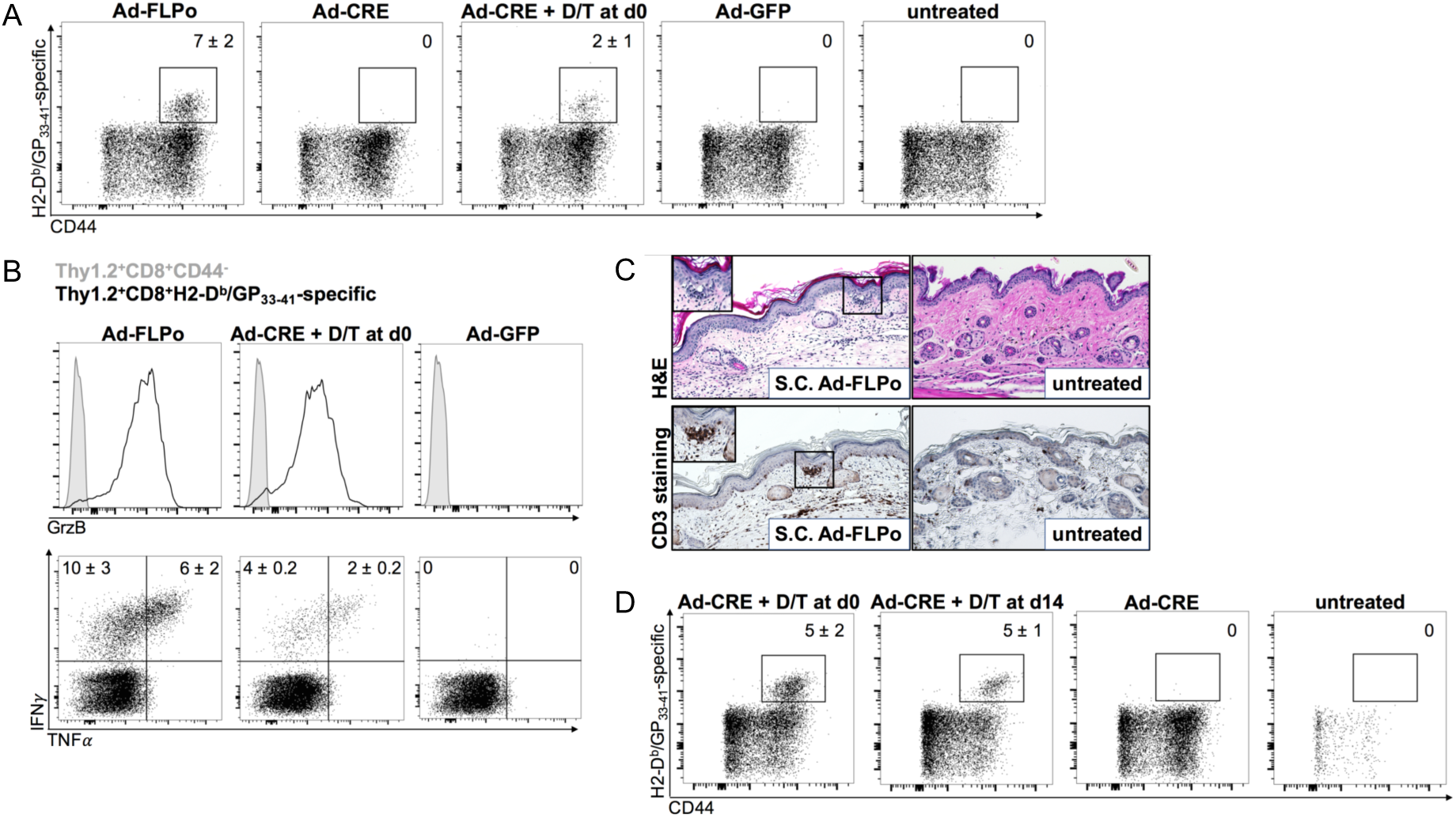
*In vivo* activation of endogenous GP_33-41_-specific T cells in NINJA mice after local infection with Ad-FLPo. **a-d,** Endogenous neoantigen-specific effector CD8 T cell responses against self antigens. NINJA x CAG-rtTA3 mice were left untreated or infected S.C. infection in the footpad with (**a-c**) Ad-FLPo, Ad-Cre (alone or administered together with doxycycline and 4-OHT), Ad-GFP, or (**d**) Ad-Cre (alone or administered either together with doxycycline and 4-OHT or 14 days prior to treatment with doxycycline and 4-OHT). **a**, FACS plots show frequency of endogenous activated (CD44^+^) H2Db/GP_33-41_-specific Thy1.2^+^CD8^+^ cells from the day 8 spleen. **b**, Histograms show Granzyme B expression (top) and FACS plots show IFNγ and TNFα production (bottom) by spleen H2Db/GP_33-41_-specific Thy1.2^+^CD8^+^ cells from mice. Representative dot plots are shown (*n* = 9). **c**, Histological sections of footpad show epidermal and dermal thickening and infiltration by endogenous CD3^+^ cells in NINJA x CAG-rtTA3 mice infected S.C. with Ad-FLPo (top and bottom left) as compared to untreated NINJA x CAG-rtTA3 mice (top and bottom right). Representative images (*n* = 3). **d**, Temporally delayed antigen induction allows for neoantigen-specific CD8 T cell responses. FACS plots show the frequency of endogenous activated (CD44^+^) H2Db/GP_33-41_-specific Thy1.2^+^CD8^+^ cells from local draining lymph nodes of NINJA x CAG-rtTA3 mice 8 days after neoantigen induction. Representative dot plots (*n* = 8). D = doxycycline; T = 4-OHT. Average values ± SEM are shown in **a**, **b** and **d**.

Next, we tested whether the regulatory module was functional *in vivo* by crossing NINJA mice to transgenic mice expressing rtTA3 under a ubiquitous promoter (CAG-rtTA3) and infecting NINJA x CAG-rtTA3 in the footpad with 10^7^ PFU Ad-Cre while treating them with D/T or vehicle control. Critically, robust endogenous GP33-41-specific CD8 and GP66-77-specific CD4 T cell responses were observed 8 days after infection only in mice that were also treated with D/T, demonstrating that the regulatory module of NINJA can be used to induce the neoantigen module *in vivo* (**Fig 5A-B** and data not shown).

Importantly, we did not observe neoantigen-specific T cell responses when NINJA x CAG-rtTA3 mice were infected with Ad-Cre but not treated with D/T, suggesting the poised regulatory module was not leaky in the context of adenoviral vector infection. However, it was possible that Ad-Cre induced a small amount of neoantigen that resulted in responses by endogenous GP33-specific CD8 T cells at levels below the limit of detection. To account for this possibility, we infected NINJA mice in the footpad with 1:9 mixtures of Ad-FLPo:Ad-GFP or Ad-FLPo:Ad-Cre. In this scenario, Ad-FLPo:Ad-GFP elicits a detectable GP33-specific CD8 T cell response, and thus we would expect that if there was leaky neoantigen expression after Ad-Cre, this would enhance the relatively weak Ad-FLPo response in Ad-FLPo:Ad-Cre infected mice. However, we did not observe increases in the magnitude of the endogenous neoantigen-specific T cell responses in Ad-FLPo:Ad-Cre-infected mice compared to Ad-FLPo:Ad-GFP-infected mice (**Fig S9**), consistent with the hypothesis that, even in the poised state, regulation of NINJA *in vivo* is maintained tightly.

### Delayed neoantigen induction triggers responses by neoantigen-specific T cells

To demonstrate the potential of NINJA for applications where dissociating infection and neoantigen induction would be preferable, we next tested a protocol where we infected NINJA x CAG-rtTA3 mice in the footpad with 10^7^ PFU Ad-Cre and then waited 14 days before treating them with D/T. Compared with Ad-Cre-infected NINJA x CAG-rtTA3 mice that were treated with D/T at the time of infection, we observed a similar neoantigen-specific endogenous GP33-specific CD8 T cell responses (**Fig 5D**), suggesting that NINJA mice can be used to trigger neoantigen-specific T cell responses against self-neoantigens under conditions that are not overtly inflammatory.

## Discussion

One of the barriers in studies of T cell immunology has been the absence of mouse models with inducible neoantigens that allow for robust endogenous neoantigen-specific CD8 and CD4 T cell responses ^3^. Here, we engineered the NINJA mouse model to bypass central and peripheral tolerance by using many levels of regulation to create neoantigens upon induction. This process involved the development of spliced neoantigen-coding genes that require FLPo-regulated inversion ^21, 22^. This structure enabled “switch-like” induction of neoantigen expression without leakiness. To allow *in vivo* control over the switch, we engineered the regulatory module of NINJA to encode FLPo. We validated that the genetic elements in this system were functional both through *in vitro* analyses, tumor cell line models, and *in viv*o in NINJA mice. Moreover, we demonstrated that endogenous neoantigen-specific T cells in NINJA mice bypass central and peripheral tolerance mechanisms and respond robustly to local expression of neoantigens by self-tissues. This also confirmed that in NINJA mice, upon *in vivo* FLPo-mediated recombination of the neoantigen module, neoantigens are expressed and functionally presented to the components of the immune system. Moreover, we found that we could temporally dissociate the activation of the regulatory module and the neoantigen module, which still resulted in neoantigen-specific CD8 T cell responses two-weeks after the initial infectious event. This would be particularly useful for genetically engineered cancer models (like the KP model) where it is not currently possible to induce neoantigen expression independently from tumor induction. Together, these data demonstrate that NINJA is a tightly regulated system for inducible neoantigen expression *in vivo* and is capable of eliciting strong endogenous T cell responses.

The thymus is an organ that is designed to generate central tolerance towards self-antigens. Thymocytes are negatively selected when they encounter low levels of self-antigens expressed by medullary thymic epithelial cells (mTECs) ^1, 2^. This alters their development by promoting death or conversion to a regulatory phenotype (like Tregs). The expression of tissue-restricted antigens is driven by the autoimmune regulator (AIRE), and AIRE deficiency leads to Autoimmune Polyendocrinopathy Syndrome type 1 (APS-1; also known as autoimmune polyendocrinopathy-candidiasis–ectodermal dystrophy/dysplasia; APECED) ^24^. Yet, it is this high sensitivity to low level expression of self-antigens that has confounded investigators when developing mice with inducible neoantigens. As a consequence, there are many questions regarding peripheral T cell tolerance and autoimmune disease that remain unanswered and NINJA should provide a critical tool by which investigators can begin to address them.

NINJA uses three levels of regulation to control neoantigen induction, but in NINJA-C mice, one of these levels is removed and this results in expression of neoantigens in <1% of cells. This highlights the critical importance of the multiple layers of regulation in NINJA. Interestingly, the pattern of leaky expression in NINJA differs from traditional inducible-models in which a low-level expression is observed from many cells (*i.e.,* all cells in a tissue or the body) ^7^. By contrast, in NINJA-C mice, a tiny fraction of cells had full expression of neoantigens, which demonstrates the consequences of leaky expression in a switch-like system. Interestingly, even though only a few cells are neoantigen-positive in NINJA-C mice, they have complete central tolerance towards the neoantigens (data not shown). This efficiency seems to reflect the underlying biology of central tolerance, as only a small fraction (<5%) of mTECs express each tissue-restricted antigen ^3^. Critically, we have found that when NINJA mice are bred to mice with Cre expressed under a tissue-specific promoter, the development of tolerance is highly dependent on promoter, as some (like K14-Cre or PDX1-Cre) lead to central tolerance while others (like Alb-Cre) do not (data not shown). Moreover, an additional layer of regulation added by using a CreER transgene instead of Cre (*i.e.,* PDX1-CreER) abrogates these central tolerance phenotypes.

As mentioned above, NINJA is a tool that will enable research in a number of areas in immunology. NINJA was designed to be compatible with many Cre-driven genetically engineered mouse cancer models and will be useful in programming tumors from different cancers to express neoantigens, and thus facilitating studies in cancer immunology. Additionally, NINJA allows for the inducible expression of neoantigens in the absence of inflammation, which will be useful for studies on peripheral tolerance and autoimmune disease. Here, a particularly useful feature of NINJA is the ability to express neoantigens in different cell types or tissue types and determine how targeting these cells causes immune-mediated disease. Along these lines, NINJA may be useful as a model for the study of immune related adverse events (irAEs) after immune checkpoint therapy. NINJA will also be useful in the field of transplantation biology, as one could potentially transplant a syngeneic organ and then express a foreign antigen in that organ after graft acceptance. Finally, the technology in NINJA can be adapted for other scenarios where spatio-temporal control of switch-like induction is desired, such as the induction of suicide genes in specific cell types. Thus, we anticipate that NINJA has many potential applications across a wide range of fields.

## Methods

All studies were carried out in accordance with procedures approved by the Institutional Animal Care and Use Committees of Yale University and the Massachusetts Institute of Technology. All mice were bred in specific pathogen-free conditions. For experiments, 6-12 week old mice were used.

### Development and testing of neoantigen module constructs

<u>Neoantigen Module Version 1 (NM.1):</u> We designed a proof-of-concept construct, in which the GP33-80 region of LCMV was fused to the C-terminus of eGFP (NM.1; **Fig S2A**). To design the DNA sequence (*in silico*) for our initial construct (see **Fig S1B** for details), we started with the fusion of eGFP to GP33-80 and then introduced a consensus splice donor (SD) sequence AAG|GTGAGT 5’ to the DNA encoding the K amino acid residues highlighted above in GP33-41 and GP66-77 (note, AAG encodes for K). Just 3’ to the DNA encoding the A and G amino acid residues in GP33-41 and GP66-77 above (respectively), we inserted a strong splice acceptor (SA) sequence from human adenovirus 5. We also inserted the intronic sequences from the chicken beta-actin (cβA) gene (cβA intron or cβAi; contained in the pCAG promoter) between the splice donor and acceptor sequences. This generated a DNA construct that was eGFP:GP33:SD:cβAi:SA:GP34-GP68:SD:cβAi:SA:GP69-80, where the SA:GP34-GP68:SD portion encoded exon 2. We then replaced this exon 2 sequence with its reverse and complement (generating eGFP:GP33:SD:cβAi:**SD:GP68-GP34:SA**:cβAi:SA:GP69-80), because we reasoned in this configuration exons 1 and 3 would splice directly. Splice sites were confirmed using the GENESCAN web server at MIT (http://hollywood.mit.edu/GENSCAN.html). Finally, to ensure no spurious transcription of exon 2 could occur, we inserted an inverted SV40 poly adenylation sequence (iSV40pA) between the 3’ FRT-F3:FRT-WT sequences.

We then synthesized the backbone DNA for the NM.1 construct containing the WT and F3 FRT sites in opposite orientations and multiple restriction sites to facilitate the cloning of the subsequent parts in the pIDTBlue vector (Integrated DNA Technologies). The vector was synthesized in two halves due to the repetitiveness of the sequence. These halves were jointly cloned into the pBluescript II SK-vector (Agilent Technologies). This construct had the orientation: GP33:SD:intron1:FRT-F3:FRT-WT:**SD:GP68-GP34:SA**:FRT-F3:iSV40pA:FRT-WT:intron2:SA:GP69-80. PCR amplification was used to clone GFP with a splice donor into the construct upstream of the GP33. This construct was then cloned into the expression vector pcDNA3 (Thermo) for testing and validation. Note, the DNA sequences for all constructs are provided as a Supplementary Sequences document.

We transiently transfected 293T cells with the NM.1 construct alone or co-transfected the construct with a pcDNA3 plasmid containing FLPo recombinase. Flow cytometric (FACS) analysis showed the construct alone was fluorescent when the neoantigen was in the OFF state (no FLPo), but recombination was induced, we noted a loss of fluorescence intensity in the cells (**Fig S2B**). To determine if the protein of the correct size was produced, protein from transfected 293T cells was analyzed by western blotting with an anti-GFP antibody (clone D5.1, Cell Signaling). This corresponded with an increase in protein size from 27 to 32 kDa by western blotting (**Fig S2C**). We also confirmed that the correctly spliced products were generated by generating cDNA from 293T cells transfected with the NM.1 construct with and without FLPo. cDNA was amplified using using PrimeSTAR HS DNA Polymerase (Takara Bio Inc., cat. R040A) and the primers GFP F AgeI 72 and GP80 R XhoI 64 (**Table S1**). This PCR fragment was cloned into the pCR-Blunt TOPO kit (ThermoFisher Scientific, cat. K280002) and DNA from 7-10 clones from before and after FLPo-mediated recombination was prepared by miniprep (Qiagen, cat. 27106) and sequenced by Sanger sequencing with the primer GFP int F Age 66. All of the clones had the correct sequences (data not shown).

<u>Version 2 (NM.2)</u>: Generated to test if shorter intron sequences would function, to reduce the chances of spurious DNA recombination in cloning for future versions.

<u>Version 3 (NM.3):</u> *Linking neoantigen expression to fluorescence.* This approach was inspired by the biofluorescence complementation (BiFC) assay ^25^. Yellow fluorescent protein (YFP) and GFP have beta-barrel structures consisting of 11 beta-strands that are connected by loops, and, for BiFC, YFP or GFP is split between amino acid residues 155 and 156 (beta-strands 7 and 8) to generate two halves. We inserted the spliced neoantigen construct from NM.2 between amino acid residues 155 and 156 of YFP (NM.3 - in frame; **Fig S3A-B**). In the OFF-state, YFP would also contain amino acids 33 and 69-80 from GP within the loop connecting beta-strands 7 and 8 of YFP (**Fig S3B**).

To generate a construct that would *become* fluorescent after FLPo induction, we next engineered a frame shift such that YFP molecule would require exon 2 to maintain the correct reading frame (*i.e.,* direct splicing of exons 1 and 3 would result in a splicing event that would alter the reading frame; **Fig S3C**). This was achieved *in silico* by adding an A residue into the splice donor sequence at the 3’ end of exon 1 and deleting a G from the splice acceptor sequence at the 5’ end of exon 2. Next, we engineered the 5’ end of exon 3 to contain a TTC -> TTT mutation that was silent when exon 2 spliced to exon 3 (ON state) but created a premature truncation when exon 1 spliced to exon 3. This would ensure that the OFF state produced only the N-terminal half of YFP.

We synthesized the NM.3 construct with the frameshift, transfected the variant into 293T cells and analyzed their fluorescence by FACS. As a control, we mutated the NM.3 construct to remove A^463^, such that the splicing of exon 1 to exon 3 would occur in frame. As expected, without the frameshift the construct was fluorescent in the OFF state (**Fig S3D**). Introduction of the frameshift into the NM.3 construct eliminated the fluorescence of YFP in the OFF state by FACS and resulted in production of only the N-terminus of YFP by Western (**Fig S2C**). However, the construct was not fluorescent when co-transfected with plasmid containing FLPo (**Fig S3D**), despite the production of a correctly sized protein (**Fig S2C**).

<u>Versions 4-6 (NM.4-NM.6)</u>: To address the lack of fluorescence, we attempted a number of modifications to the construct including the addition of linkers between the N- and C-terminal halves of YFP and GP33-80 (NM.4) and incorporating GFP instead of YFP to increase the sensitivity of detection by FACS (NM.5). These did not increase the fluorescence of the constructs in the ON state by FACS, despite Western and immunofluorescence (IF) data showing the constructs were producing the correct proteins (**Fig S2B-C and S3E**).

We noted that by IF the construct appeared to become less sharp after FLPo and we reasoned this could indicate a protein folding error. Further investigation of the GP33-80 sequence revealed that it contained a hydrophobic region between GP43-GP59, that was strongly predicted to be a transmembrane domain (**Fig S3F**; ^26^). To address whether this hydrophobic region was preventing fluorescence, we replaced the sequence with a FLAG tag sequence, which eliminated the transmembrane domain (**Fig S3F**; NM.6). The resulting construct was indeed fluorescent in the ON state (**Fig S2D**). Thus, we had developed an inversion-mediated fluorescent neoantigen that was regulated by FLPo.

<u>Version 7 (NM.7):</u> *Neoantigen non-immunogenic in the OFF state.* In the unlikely event there was transcription of Exon 2 (in the opposite direction), there was a concern that a peptide could be produced that could mimic GP33-41 and cause tolerance. Note, there is an upstream SV40 poly A sequence in the inverted direction to minimize this risk. We analyzed the potential peptide products from this construct and realized that, due to the structure of the splice acceptor, the DNA sequence encodes GP34-41 and that the amino acid proceeding this was encoded by nCA, which could encode an Ala, Ser, Thr, or Pro residue (**Fig S4A**). We calculated the predicted binding of each peptide (GP34-41 or GP33-41 with K33A, K33S, K33T, or K33P mutations, **Fig S4B**) and found that K33A, K33S, and K33T bound as strongly or more strongly to H2-D^b^ as the wt GP33-41 peptide. We also confirmed that K33A, K33S, and K33T bind to H2-D^b^ using RMA-S cells, and that K33A and K33S could activate GP33-specific P14 TCR Tg CD8 T cells (**Fig S4C** and data not shown). By contrast GP34-41 and K33P did not bind to H2-D^b^ or stimulate T cell activation, although GP34-41 did bind to H2-K^b^ as previously reported^27^. We engineered the splice acceptor in Exon 2 so that it encoded K33P in the OFF state and we added the amino acids GP42-43 (**Fig S2A** – NM.7), which increase the affinity of GP33-41 for D^b^ to a more physiologic level ^27^.

### Development and testing of regulatory module constructs

Development of FLPoER^251^: We generated a FLPoER fusion by fusing amino acids 282-596 of the human estrogen receptor (hER) containing the T2 mutations (G400V, M543A, and L544A; ^28^) to the C-terminus of FLPo but identified that this had relatively weak activity in reporter cells upon 4-hydroxytamoxifen treatment (4-OHT; **Fig S5A**). Previous work in yeast suggested that fusing a longer portion of ER (251-596) to FLPe recombinase can increase activity, although this was not tested in mammalian cells ^21^. We synthesized this version (henceforth called FLPoER^251^) and verified that it was more responsive to 4-OHT treatment and that its activity was not increased by estrogen (E2) treatment (**Fig S5A**). FLPoER^251^ was leakier than FLPoER, but because it was one of three elements in the regulatory module, we chose to trade-off increased leakiness for increased responsiveness to 4-OHT and proceeded with FLPoER^251^.

Creating spliced FLPoER^251^ for the regulatory module: Next, we assembled a *Rosa26* targeting vector, which consisted of pTRE:Lox-STOP-Lox (LSL):FLPoER^251^ (**Fig S5B**), 2x CGG insulator, and the neoantigen module. Upon assembly, however, we discovered that the neoantigen module in this construct was being recombined by FLPo activity in *E coli* (**Fig S5C**; note neither the LSL nor the ER^251^ prevented FLPo activity in bacteria). Thus, we needed to redesign the regulatory module so that FLPo was not expressed in bacteria. We considered introducing an intron into the sequence of FLPoER^251 29^, as this would have prevented FLPo activity in bacteria. However, we became concerned that low-level expression of FLPoER^251^ (through the LSL site) could lead to leaky activation of the neoantigen in future applications. Therefore, we designed an approach where FLPoER^251^ also required recombination before it could produce a functional protein (**Fig 1A**).

To generate the inversion-inducible FLPoER^251^ construct, we first split FLPo into N- and C-terminal halves. Based on its crystal structure, we decided to split FLPo between a-helices I and J, which would split the catalytic residues across two exons. We introduced splicing sites as before, taking advantage of K285 and D286 and engineering an intron with SA and SD sequences from intron 2 of mouse p53. Here, exon 2 was inverted relative to exon 1 and placed between non-compatible lox sites (responsive to Cre-recombinase; ^30^). Downstream of exon 2, we included a poly A sequence from herpes simplex virus (HSV), which is predicted to terminate transcription in both the 5’ -> 3’ and 3’ -> 5’ directions. Thus, in the OFF state, the HSV poly A was oriented in such a manner that it should terminate the RNA transcript for N-terminal FLPoER^251^, while in the ON state it would terminate the RNA transcript for of the full-length FLPoER^251^.

For integration into the larger construct, we reoriented the regulatory module such that it would run 3’ to 5’ relative to the neoantigen module. This meant that the pTRE-tight promoter would be next to the 2x CGG insulator and the Rosa promoter (outside of the construct) would be transcribed in the opposite direction of the regulatory module. To prevent issues with transcription in both directions and to allow for selection of clones after integration, we inserted a cassette for puromycin resistance in the 5’ most end of the regulatory module (**Fig S6**). This consisted of the PGK promoter, the puromycin resistance gene and the HSV poly A sequence. In the OFF configuration, this element would provide puromycin resistance and also terminate the endogenous transcript from the *Rosa 26* promoter. Moreover, after Cre-mediated recombination in the regulatory module, this puro resistance cassette would be genetically deleted.

### Assembly of the NINJA targeting construct

The regulatory and neoantigen modules were inserted into the pzDonor Rosa26 targeting construct separated by the 2xCGG insulator to minimize the impact of the CMV enhancer on the pTRE-tight promoter. Moreover, a pgk-puromycin resistance cassette was inserted into the regulatory module in such a way that the HSV poly A sequence was predicted to terminate transcription downstream of the Rosa26 promoter. However, the puromycin resistance cassette is lost following Cre-mediated recombination, providing an assay for recombination at this allele.

### Transient transfection of 293T cells for DNA isolation, in vitro induction of NINJA, Western Blot and IF

293T cells were cultured in complete DMEM (10% HI-FBS, 1x Pen/Strep) and then transfected using the LipoD293 Transfection reagent (SignaGen Laboratories, cat. SL100668) according to manufacturer’s instructions.

For DNA isolation, cells were transfected and harvested 48hr later and digested at 55°C overnight in lysis buffer (0.2M Tris HCl, 0.15M NaCl, 2mM EDTA, 1% SDS, 50μg/mL Proteinase K in nuclease-free H_2_O), then quenched with saturated NaCl. DNA was precipitated by incubating the supernatant with 100% EtOH, washed in 70% EtOH, dried and resuspended in nuclease-free H_2_O.

For *in vitro* induction of NINJA, 293T cells were transfected with 1μg NM.7 construct + 0.3μg of Cre, Cre/rtTA, rtTA, or FlpO construct. 24hr later, 1μg/ml of doxycycline (Millipore Sigma, cat. D9891) and 1μg/ml of 4-OHT (Millipore Sigma, cat. SML1666) were added to the culture media. Cells were harvested 72hr later to measure GFP expression by flow cytometry on a BD Accuri C6 analyzer (BD Biosciences). For Western Blot, 293T cells were transfected with 0.5ug/ml target construct + 0.15ug/ml FlpO construct. Total proteins were harvested after 72 hours using RIPA lysis reagent protocol (ThermoFisher Scientific, cat. 89900). Protein content was quantified by BCA assay (ThermoFisher Scientific, cat. 23225), and 3-5ug total proteins were mixed in 1x LDS Sample Buffer + 1x sample reducing reagent, loaded on BOLT or Nupage 4-12% BisTris precast gel (ThermoFisher Scientific, cat. NW04122BOX) and run with SeeBlue2 Protein Ladder (ThermoFisher Scientific, cat. LC5925). Western Blotting was performed according to Invitrogen’s MiniBlot Western System procedures (with NuPage MES buffers). Briefly, anti-GFP (N-terminus) (clone D5.1, Cell Signaling Technology) or anti-Grp94 (loading control) (Cell Signaling Technology, cat. 2104) primary antibodies were left overnight at 4C in 10% Milk in TBST. HRP-conjugated goat anti-rabbit secondary antibody (ThermoFisher Scientific, cat. 656120) was incubated for 1 hour at room temperature. Blot was imaged using Pierce ECL (ThermoFisher Scientific, cat. 32106) or Pierce SuperSignal West Femto ECL kits (ThermoFisher Scientific, cat. 34096).

For IF, 293T cells were transfected with 1.5μg of the indicated construct (NINJA neoantigen module NM.5 or NM.7 with neoantigen either ON or OFF) and 72hr later were fixed in 2% PFA for 20 minutes. Staining was performed as described previously^31^ using a primary antibody specific for a linear epitope of the N-term portion of GFP (clone D5.1, Cell Signaling Technology) or for a conformational epitope of GFP to detect correctly folded GFP (clone 3E6, Invitrogen).

### Lymphocyte isolation from spleen and LN

Organs were processed as described previously^32^.

To purify CD8^+^ T cells from the spleen of TCR transgenic P14 mice, the EasySep Kit (StemCell Technologies, cat. 18000) was used according to manufacturer’s instructions.

### DC2.4 proliferation and killing assay

DC2.4 cells were transfected with 1ug of target construct using the Lipofectamine3000 transfection kit (Invitrogen, cat. L3000008) following manufacturer’s instructions. 24hrs later, transfected DC2.4 cells were cocultured for 5 days with 2 * 10^6^ Thy1.1^+^/1.1^+^ P14 cells purified as detailed above and stained with CellTrace Violet Proliferation kit (Invitrogen, cat. C34557) according to manufacturer’s instructions. Killing of DC2.4 was then assessed by Crystal violet staining (0.1% Crystal Violet solution), while P14 T cells were stained with antibodies specific for Thy1.1 (clone OX-7, BioLegend), CD8 (clone 53-6.7, BioLegend), CD44 (clone IM7, BioLegend) and Vα2 (clone B20.1, BD Biosciences) and analysed by flow cytometry on a BD Biosciences LSRII analyser.

### KP-C4A3D6 cell transplant and *in vivo* P14 cell activation

fLuc^+^ P14 cells were harvested from fLuc^+^ TCR transgenic mice as detailed above, and adoptively transferred I.V. retro-orbitally into wild-type C57BL/6 (B6) recipients (5 * 10^6^ fLuc^+^ P14 cells/mouse) one day prior to tumor transplant. 2.5 * 10^4^ - 2 * 10^5^ sorted GFP^+^ NINJA-expressing KP-C4A3D6 cells (ON) or parental NINJA-negative KP-C4A3D6 (OFF) cells were transplanted into the leg muscle following baseline leg diameter measurement.

Tumor size was measured throughout the experimental time with a digital caliper (Thomas Scientific, cat. 1235C55) on two axes (diameter of coronal and sagittal plane of injected leg). Baseline measurements were taken prior to transplant, after shaving, and tumor sizes were adjusted for original leg volume. Tumor volume was calculated as (coronal diameter mm)*(sagittal diameter mm)*(average of both measurements)= mm^3^. At the end of the experimental time, tumors were harvested and dissociated in Collagenase IV (Worthington Biochemical, cat. LS004189) Buffer (1x HEPES buffer, 0.5mg/mL Collagenase IV, 20μg/mL DNase in 1x HBSS with MgCl_2_ and CaCl_2_) running the default Lung_01 protocol on a gentleMACS Dissociator instrument (Miltenyi Biotec). Tumor samples were then incubated at 37°C for 40min (rotating) prior to running the default Lung_02 protocol on the gentleMACS Dissociator. Digestion was quenched by adding 500μL FBS. Tumor samples were then strained through 70μm cell strainers, washed with 1% HI-FBS RPMI-1640 and red blood cells were lysed using 1x RBC Lysis Buffer (eBioscience, cat. 00-4333-57)

### BMDC Transplant

Thy1.1^+^/1.2^+^ P14 cells and Thy1.1^+^/1.1^+^ SMARTA cells were isolated from TCR transgenic mice as detailed above. 10^5^ P14 cells and 10^5^ SMARTA cells were co-transferred I.V. by retro-orbital injection into wild-type B6 recipients 0-19 days prior to BMDC transplant.

BMDCs were prepared from the bone marrow of NINJA mice as described in ^31^. Four days after bone marrow harvest, maturing BMDCs were transduced *in vitro* with 10^10^ PFU/mL of recombinant Ad5CMVFLPo (Iowa Vector Core, VVC-U of Iowa-530HT) or 3.5 x 10^8^ PFU/mL of recombinant Ad5CMVeGFP (Iowa Vector Core, VVC-U of Iowa-4). BMDCs were harvested 3 days later and stained with antibodies specific for CD11c (clone N418, BioLegend) and MHC-I (H-2Kb, clone AF6-88.5.5.3, eBioscience) and analysed by flow cytometry on a BD LSRII analyser (BD Biosciences). 10^4^ CD11c^+^MHC-I^+^GFP^+^ BMDCs were then transplanted into the footpad of B6 hosts in 15μl PBS.

Recipient animals were euthanized 7 days after BMDC transplant to collect spleen and the draining LN (popliteal). Organs were processed as described in ^32^ and samples were stained with antibodies specific for Thy1.1 (clone OX-7, BioLegend), Thy1.2 (clone 30-H12, BioLegend), CD4 (clone GK1.5, BioLegend) and CD8 (clone 53-6.7, BioLegend). Cells were then analysed on a BD LSRII flow cytometer (BD Biosciences).

### Creation of the NINJA transgenic lung tumor cell line

A B6 cell line derived from an Ad-CRE-infected Kras^G12D^ p53^fl/fl^ mouse was co-transfected with the NINJA targeting construct and RNA encoding a zinc-finger nuclease targeting the *Rosa26 locus* (Millipore Sigma) using Amaxa nucleofector (Lonza). We generated a KP-NINJA line and verified that the NINJA construct was targeted to the *Rosa26* locus. Single-cell clones were selected based on Puro resistance and screened for incorporation of the NINJA targeting vector by PCR. The C4 clone was selected for further study. rtTA was stably introduced into C4 cells via a dual promoter retrovirus expressing rtTA and a blasticidin resistance cassette (C4A3). A blasticidin-resistant single cell clone (C4A3D6) was selected and infected with Ad-CRE to mediate recombination of the regulatory module. Activation of the neoantigen module was achieved by incubating C4A3D6 cells in vitro with doxycycline (1ug/ml) and 4-hydroxytamoxifen (1ug/ml) for 72 hours.

### Generation of the NINJA mouse

Targeting was performed by the Swanson Biotechnology Center Preclinical Modeling facility of MIT. Briefly, NINJA mice were generated by transfecting the PvuI-HF linearized NINJA targeting construct into B6 ES cells (JM8) along with mRNA encoding ZFNs for the *Rosa26* locus (Millipore Sigma). We screened clones for proper targeting by southern blotting and identified one correctly targeted clone (**Fig S7**). We injected these targeted ES cells into B6 blastocysts and selected a high percentage chimera (>80%) for breeding to a B6 female. This breeding resulted in germline transmission in two pups that were used to produce the NINJA mouse line.

NINJA-F and NINJA-C mice were generated by crossing NINJA mice to FLPe or Cre deleter mice (JAX, stock # 003946) and (JAX, stock # 006054) respectively and then crossing FLPe+ or Cre+ F1 mice to B6 mice and screening the progeny for FLPe- or Cre-driven recombination at the NINJA locus. Mice were bred to homozygosity and maintained as independent strains.

NINJA x CAG-rtTA3 were generated by crossing NINJA mice to CAG-rtTA3 mice (JAX, stock # 016532).

### Genotyping of NINJA, NINJA x CAG-rtTA3, NINJA-C, and NINJA-F mice

Mouse tail DNA was amplified using KAPA Taq PCR Kit (Takara Bio, cat. R040A) and primers Rosa26 WT F, Rosa26 WT R, and Rosa26 Ninja R (**Table S1**). Expected bands were 378bp for WT allele and 280 bp for NINJA allele. For CAG-rtTA3 genotyping, primers CAGrtTA common F, CAGrtTA WT R and CAGrtTA Transgene R (**Table S1**) were used and expected bands were 360bp for WT allele and 330bp for Transgene. For NINJA-C, Rosa26 WT F, Rosa26 NINJA R, and Rosa26 Ninja-C R primers (**Table S1**) were used for expected bands of 280 bp for unrecombined NINJA allele or 694 bp for NINJA allele with recombination in RM. For NINJA-F mice, Rosa26 Ninja-F F and Rosa26 Ninja-F R primers (**Table S1**) were used for expected bands of 784 bp for unrecombined NINJA or 442 bp for NINJA allele with recombination in RM.

### Southern Blot

DNA from ES cell clones was digested with BamHI or EcoRV overnight and then was run on a 0.7% TAE gel overnight at 25-35 V. Gel was HCl depurinated, NaOH denatured, and neutralized using standard protocols and transferred passively to Hybond XL membrane via wicking method. Blot was UV-crosslinked using a Stratalinker (Stratagene) and was probed using dCTP-32P labeled probes made using the PrimeIT II kit (Agilent, cat. 300385) and Roche Quick Spin Columns (TE, cat. 11532015001). The template for the southern probe was made from a 817 bp fragment amplified from B6 genomic DNA with primers Rosa26 southern probe F and R (**Table S1**, ^33^).

### P14/SMARTA crosses

NINJA, NINJA-C and NINJA-F mice were crossed with either GP33-43-specific TCR transgenic P14 mice or with GP66-77-specific TCR transgenic SMARTA mice. Developing T cells were harvested from the thymus of 6-8 weeks old litters and analysed by flow cytometry on a BD Biosciences LSRII analyser after staining with antibodies specific for Thy1.2 (clone 30-H12, BioLegend), Thy1.1 (clone OX-7, BioLegend), CD8 (clone 53-6.7, BioLegend), CD4 (clone GK1.5, BioLegend).

### Analysis of PBMCs for NINJA expression

6-10 weeks old NINJA, NINJA-C, NINJA-F and wild-type B6 mice were bled retro-orbitally using heparinized glass capillaries (Fisher Scientific, cat. 22362566) to collect 50-100uL of blood into 0.04% Citrate Buffer. Red blood cells were lysed using 1x RBC Lysis Buffer (eBioscience, cat. 00-4333-57) and the remaining PBMCs were analysed by flow cytometry on a BD Biosciences LSRII analyser for detection of GFP expression.

### LCMV-Armstrong infections

7-10 weeks old NINJA, NINJA-F or age-matched wild-type B6 mice were infected intraperitoneally with 2 * 10^5^ PFU/mouse of LCMV-Armstrong. For P14 tolerance induction assay, Thy1.1^+^/1.1^+^ 10^6^ P14 cells were adoptively transferred through retro-orbital I.V. injection into recipient mice two weeks before infection with LCMV-Armstrong. Mice were euthanized 8 days after infection to collect and process spleens as described in ^32^.

### In vitro antigen-specific restimulation and flow cytometry

For antigen-specific restimulation, 1 * 10^6^ splenocytes were cultured for 6hr at 37°C in 5% CO_2_ in complete RPMI-1640 (10% HI-FBS, 55μM beta-mercaptoethanol, 1x Pen/Strep and 1x L-Glut) supplemented with LCMV GP33-41 peptide (0.5μg/mL, AnaSpec, cat. AS-61296) or left unstimulated.

For flow cytometric analysis, 1 * 10^6^ splenocytes were stained with antibodies specific for surface markers Thy1.2 (clone 30-H12, BioLegend), Thy1.1 (clone OX-7, BioLegend), CD8 (clone 53-6.7, BioLegend), CD4 (clone GK1.5, BioLegend), CD44 (clone IM7, BioLegend) and tetramers for either H2Db/GP_33-41_-specific CD8 T cells, H2Db/NP_396-404_-specific CD8 T cells or I-A^b^/GP_66-77_-specific CD4 T cells (all from NIH Tetramer Core Facility). For intracellular staining, cells were processed using the Fixation/Permeabilization kit from BD Biosciences (BD Cytofix/Cytoperm, cat. 554714) following manufacturer’s instructions and stained with antibodies specific for IFNγ (clone XMG1.2, BD Biosciences) and TNFα (clone MP6-XT22, eBioscience). Samples were analysed on a BD Biosciences LSRII flow cytometer.

### Adenoviral vector infections and in vivo induction of NINJA

7-10 weeks old NINJA or NINJA x CAG-rtTA3 mice or age-matched wild-type B6 mice were infected with a dose of 10^7^ PFU/mouse of recombinant Ad5CMVFLPo (Iowa Vector Core, VVC-U of Iowa-530HT), Ad5CMVCre (Iowa Vector Core, VVC-U of Iowa-5-HT) or Ad5CMVeGFP (Iowa Vector Core, VVC-U of Iowa-5-HT). Adeno vectors were either injected retro-orbitally (I.V.), intratracheally (I.T.), intramuscularly (I.M.) or subcutaneously in the footpad (S.C.), depending on the experiment.

For induction of neoantigen with Ad5CMVCre infection, mice were fed doxycycline-containing food (Envigo Teklad, cat. TD.120769) for four consecutive days starting one day prior to S.C. infection with Ad5CMVCre. Mice also received two consecutive doses of a 50mg/mL solution of 4-hydroxytamoxifen (Millipore Sigma, cat. T176) in DMSO (AmericanBIO, cat. AB03091) that was painted on the surface of the infected footpad (10-20uL/footpad) immediately following infection with Ad5CMVCre and the day after infection.

### *In vivo* imaging by IVIS

For *in vivo* imaging of mice implanted with tumor cells, equal numbers (from 2.5 x 10^5^ - 2 x 10^6^) GFP^+^ and GFP^-^ KP-NINJA cells were implanted intramuscularly (I.M.) in opposite legs of shaved B6 mice (1 to 14 days before), and adoptively transferred with 10^4^ fLuc^+^ P14 cells by retro-orbital injection (I.V.). Mice were imaged on IVIS as detailed below on days 2, 7, 9, and 11 or 12 after tumor transplant. Tumors were measured every other day as detailed above.

For *in vivo* imaging of mice following adenoviral vector infection, mice were adoptively transferred with 5 x 10^3^ fLuc^+^ P14 cells by retro-orbital injection (I.V.) one day prior to infection with Ad5CMVFLPo. Eight days after infection, hair was removed and mice were injected I.V. with a PBS solution containing 3mg of luciferin/mouse (XenoLight D-Luciferin - K+ Salt, PerkinElmer, cat. 122799) and imaged using an IVIS Spectrum *In Vivo* Imaging system (PerkinElmer) instrument.

### IF/IHC

For IF/IHC, mice were adoptively transferred with 5 x 10^3^ dsRED^+^ P14 cells by retro-orbital injection (I.V.) one day prior to S.C. infection with Ad5CMVFLPo in the footpad. Eight days after infection, mice were euthanized to collect infected footpads and the untreated contralateral footpads (negative control). For IF, tissues were fixed overnight at 4°C with a paraformaldehyde-lysine-periodate fixative (PLP), subsequently cryoprotected with 30% sucrose in PBS for 6-8hr at 4°C, and then embedded in cryomolds with 100% optimum cutting temperature (O.C.T.) compound (VWR, cat. 25608-930) on dry ice for freezing.

For IHC, tissues were fixed in a 1x formaldehyde solution in PBS (Millipore-Sigma) and then embedded in paraffin and sectioned by the Yale Pathology Core Facility. All tissues were sectioned and stained as previously described^31^.

## Acknowledgements

We thank Joshi and Jacks lab members for reviewing the manuscript and E. Sun for helpful discussion. We also thank the Yale Cancer Center (P30 CA016359 40), Yale Flow Cytometry Core, Yale School of Medicine Histology Facility, Dr. Peter Cresswell for confocal microscopy, Swanson Biotechnology Center Preclinical Modeling facility of MIT for ES cell targeting. This work was supported by grants from the Howard Hughes Medical Institute (T.J.), the K22 transition to Independence grant NCI-K22CA200912 (N.S.J.), the Damon Runyon Cancer Foundation (N.S.J.), The G. Harold & Leila Y. Mathers Foundation (N.S.J), and the National Institute of Diabetes and Digestive and Kidney Diseases of the National Institutes of Health under Award Number P30KD034989 (N.S.J.). The content is solely the responsibility of the authors and does not necessarily represent the official views of the National Institutes of Health. T.J. is a Howard Hughes Investigator and a Daniel K. Ludwig Scholar. Figures created with Biorender.com.

## Author Contributions

M.D., B.F. and N.S.J. designed studies. N.S.J. and T.J. supervised research. M.D., B.F., Y.L, M.N., I.W., J.C., K.A.C., Y.F., E.A-G., D.L., G.P.C., V.G., L.S., A.B., J.H.W., C.C., P.G., P.C., and N.S.J. conducted research

---

**Table S1.**
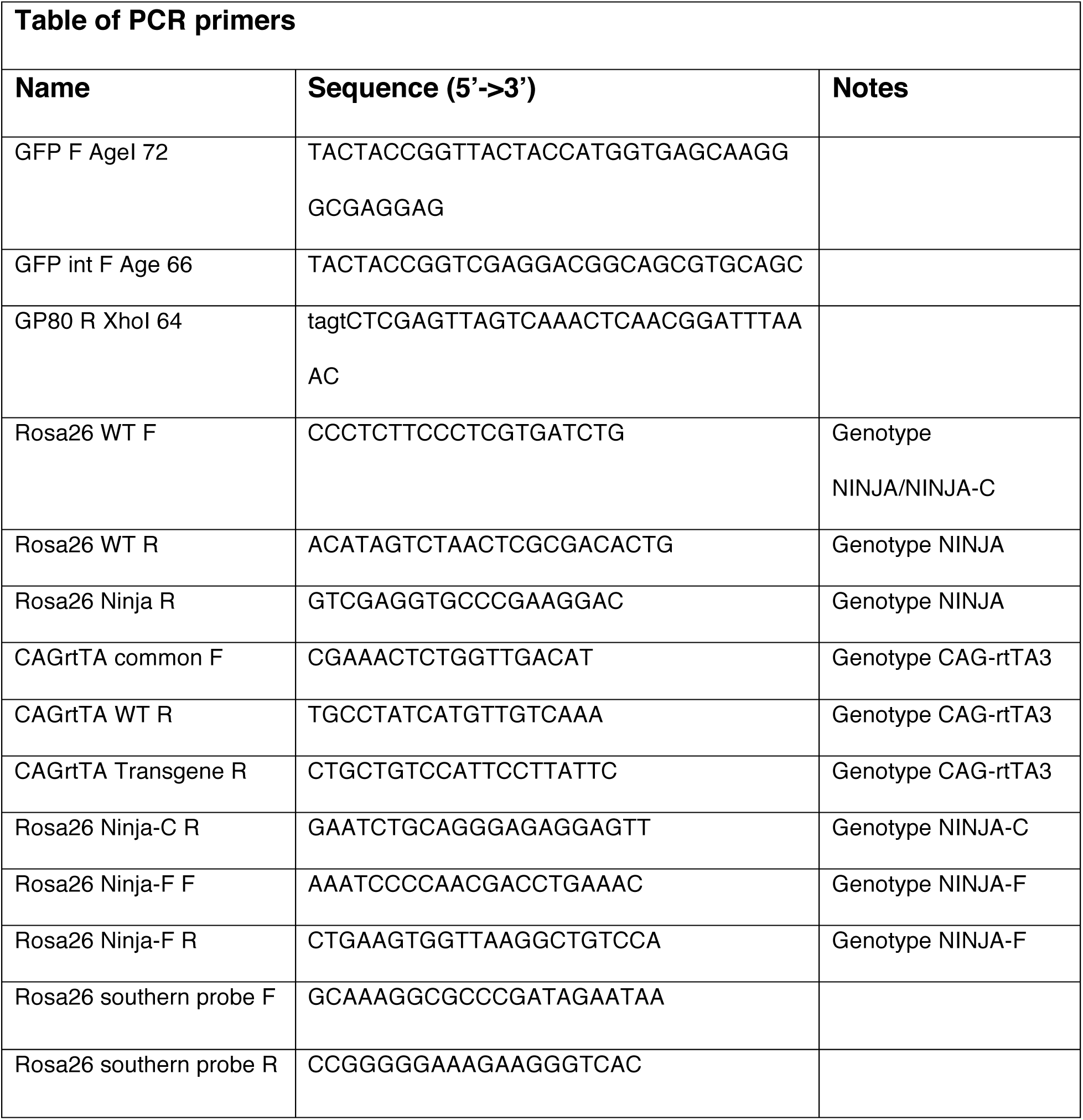

