## Supplementary materials for "NINJA: an inducible genetic model for creating neoantigens *in vivo*"

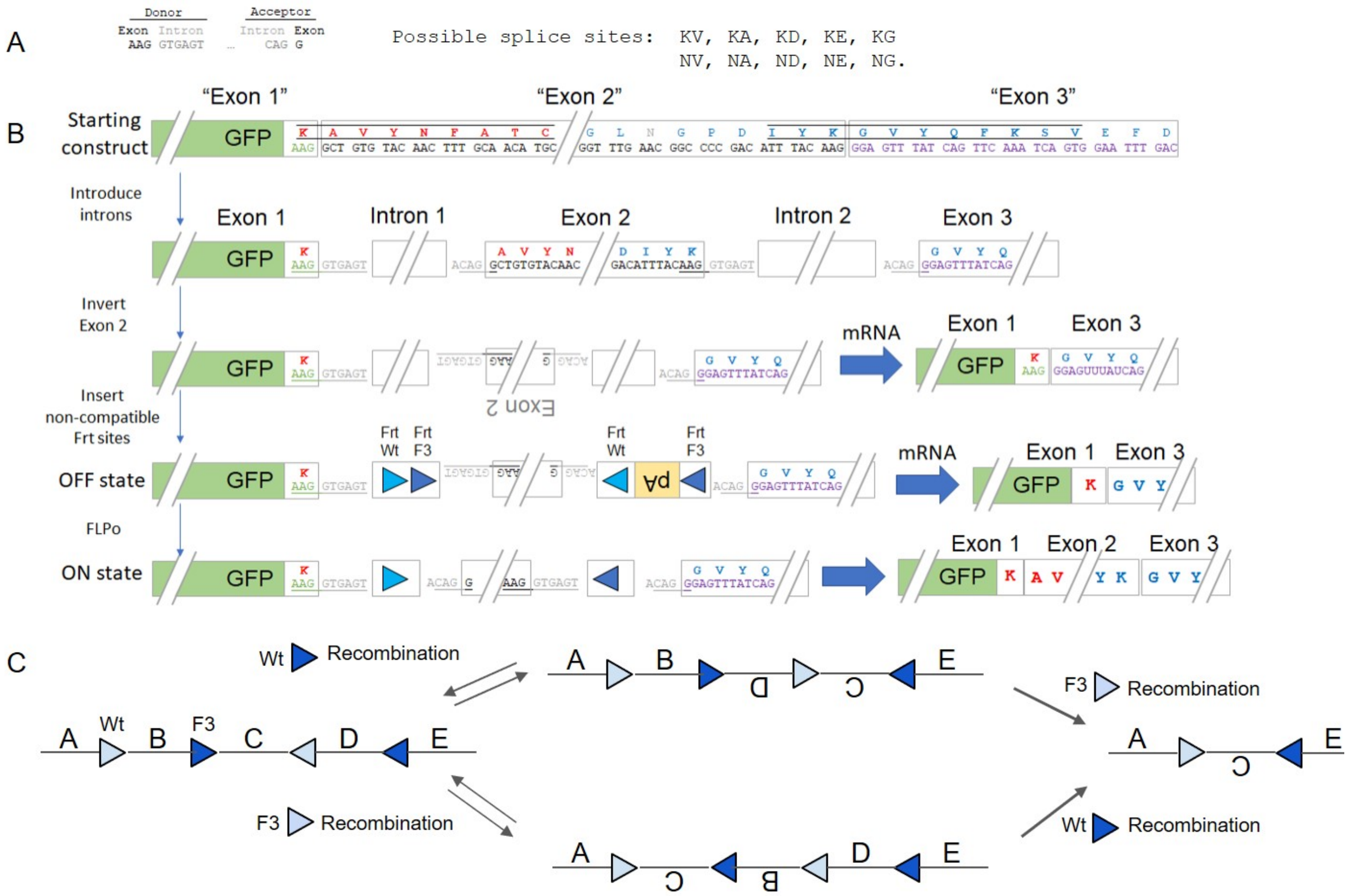


*Figure S1 - Introduction of splice sites into neoantigen module*

**a**, Possible splice donor and acceptor sequence candidates in the neoantigen module to create Exon 2. **b**, Design and inversion of Exon 2, with transcription in OFF state resulting in splicing directly from Exon 1 to Exon 3. Insertion of noncompatible Frt sites shown in light and dark blue arrows. **c**, FLPo recombinase activity results in permanent inversion and transcription of all exons of the neoantigen module.

A

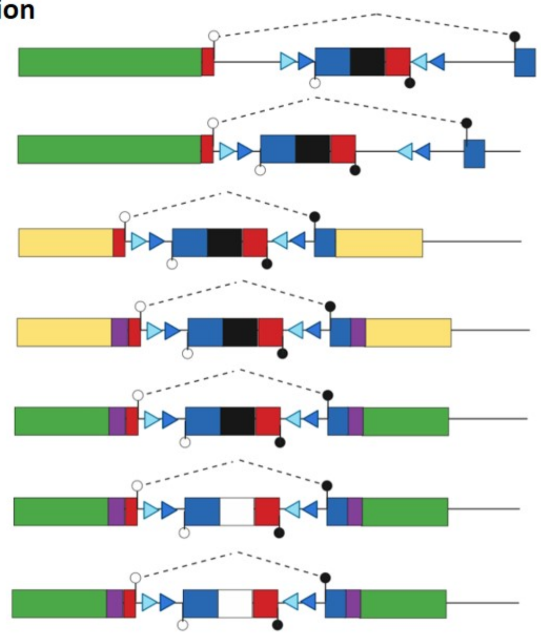

| NM Version | Modification from previous version | Antigen OFF | Antigen ON |
|---|---|---|---|
| NM.1 | Proof of concept | Fluorescent | Fluorescent |
| NM.2 | Introns 1 and 2 were identical. Replaced with short intron 1 | Fluorescent | Fluorescent |
| NM.3 | Proof of concept for split YFP design | Not Fluorescent | Not Fluorescent |
| NM.4 | Added linkers to enhance fluorescence | Not Fluorescent | Not Fluorescent |
| NM.5 | Switched to GFP to enhance detection of fluorescence | Not Fluorescent | Not Fluorescent |
| NM.6 | Replaced GP42-59 with FLAG epitope | Not Fluorescent | Fluorescent |
| NM.7 | Added back GP42-43, modified splice site in OFF state, added in frame stop in OFF state | Not Fluorescent | Fluorescent |

B

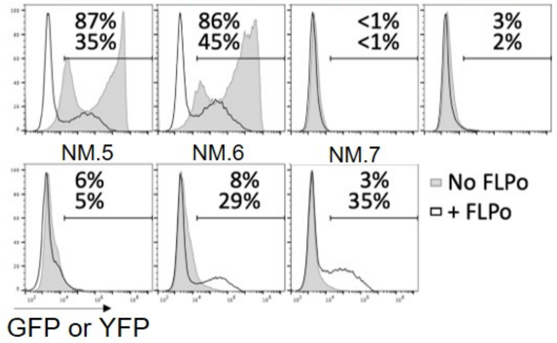


C

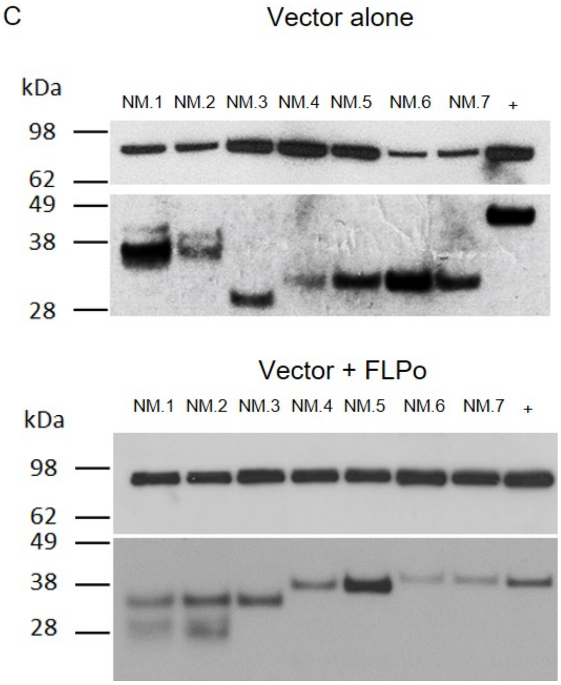

***Figure S2 - Development of advanced versions of neoantigen module***

**a**, Schematic shows 7 versions of the neoantigen module, with modifications made at each step and the fluorescence status at either ON or OFF state. **b**, Flow cytometry histograms of GFP or YFP fluorescence in each version, with or without FLPo activation. **c**, Western blotting for GRP94 (control, top panel) or N-terminal GFP (bottom panel) on lysates from 293T cells transiently transfected with each version of neoantigen module, with (top blot) or without (bottom blot) FLPo. Positive control (+) is cell lysate from KP-C4A3D6 after FLPo.

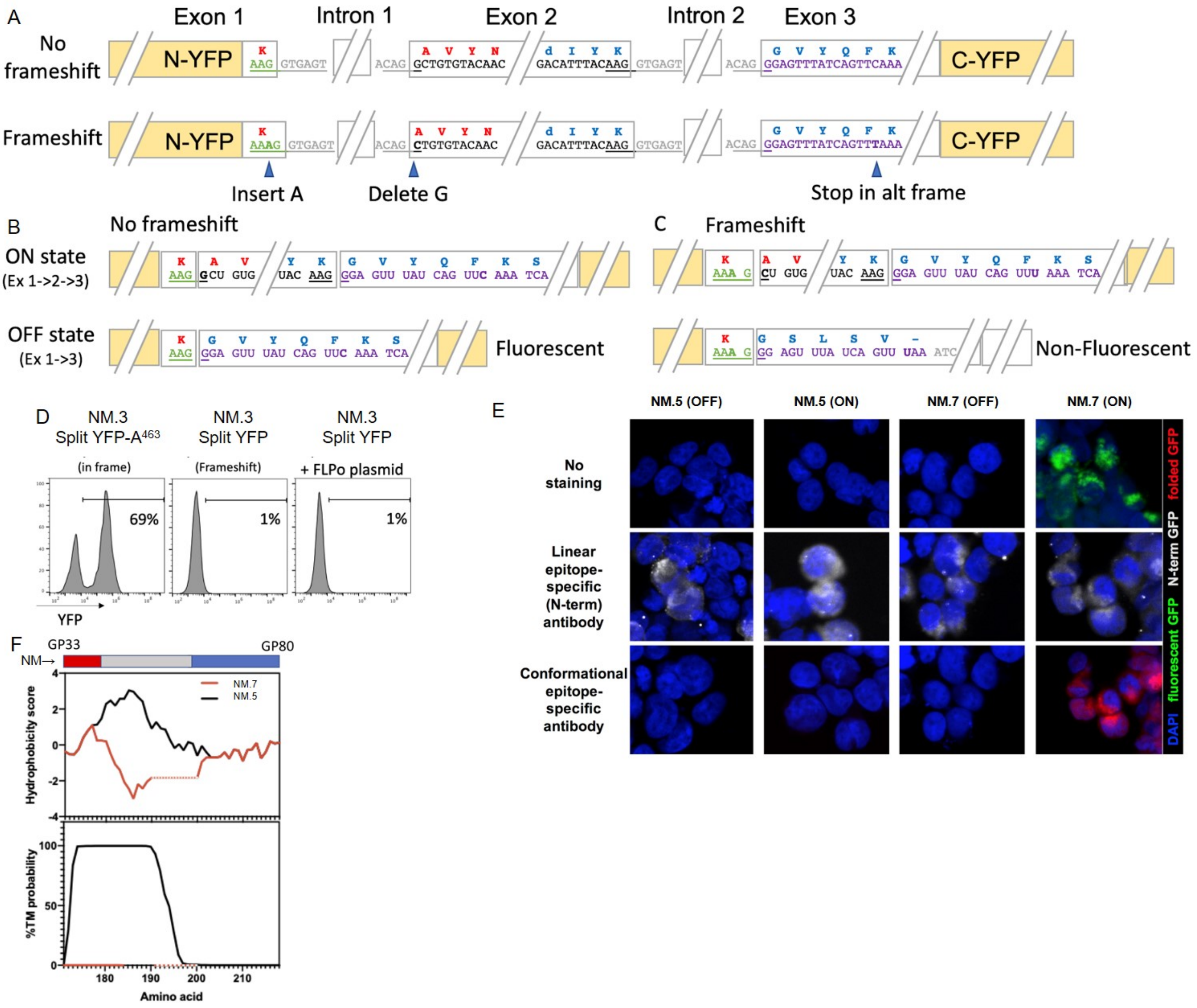

A
Exon 1
Intron 1
Exon 2
Intron 2
Exon 3
No frameshift
N-YFP
C-YFP
Frameshift
Insert A
Delete G
Stop in alt frame
B
No frameshift
ON state (Ex 1->2->3)
OFF state (Ex 1->3)
Fluorescent
C
Frameshift
Non-Fluorescent
D
NM.3 Split YFP-A463 (in frame)
69%
NM.3 Split YFP (Frameshift)
1%
NM.3 Split YFP + FLPo plasmid
1%
YFP
E
NM.5 (OFF)
NM.5 (ON)
NM.7 (OFF)
NM.7 (ON)
No staining
Linear epitope-specific (N-term) antibody
Conformational epitope-specific antibody
folded GFP
N-term GFP
fluorescent GFP
DAPI
F
GP33
GP80
NM→
NM.7
NM.5
Hydrophobicity score
%TM probability
Amino acid

***Figure S3 - Introduction of frameshift into neoantigen module***

Schematic of NM.3, which shows the two possible *in silico* insertions of the spliced neoantigen construct from NM.2 in YFP, where in one skipping exon 2 results in a frameshift and premature stop codon. **b-c**,The transcription product and fluorescence status in the ON or OFF state is shown for the frameshift version (b) or no frameshift version (c). **d**, Flow cytometry histograms of YFP fluorescence of the constructs from ©. The frameshift version is only YFP positive after FLPo exposure. **e**, 293T cells transiently transfected with plasmids expressing the indicated version of the NINJA neoantigen module (NM) were imaged by confocal microscopy after staining with an antibody specific for either the N-term portion of GFP (middle panels), for a conformational epitope of GFP (bottom panels) or with no antibody (top panels). BLUE = DAPI, RED = folded GFP, GREEN = fluorescent GFP. Representative images are shown ($n$ = 3). **f**, Hydrophobicity score (top graph) in relation to amino acid position along the neoantigen module (red/grey/blue rectangle). In version NM.5 (black line) positions GP43-GP59 are predicted to be a transmembrane domain (bottom graph), and this elevated hydrophobicity is abrogated when replaced by a FLAG domain in NM.7 (red line).

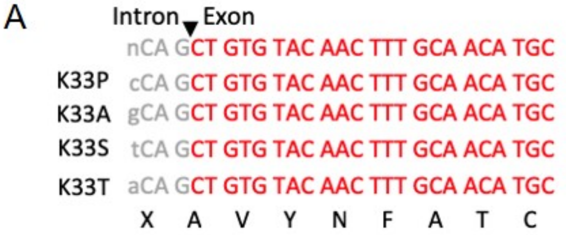


B

| Sequence | Binding affinity IC50 (nM) H-2 Db | Binding affinity IC50 (nM) H-2 Kb |
|---|---|---|
| SIINFEKL | 3220.9 | 19.4 |
| SAVYNFATC | 843.2 | 696.0 |
| AAVYNFATC | 1412.7 | 1623.3 |
| KAVYNFATC | **2444.7** | 2151.5 |
| TAVYNFATC | 3008.3 | 1317.2 |
| _AVYNFATC | 24917.4 | 29.9 |
| PAVYNFATC | 20007.9 | 11264.5 |

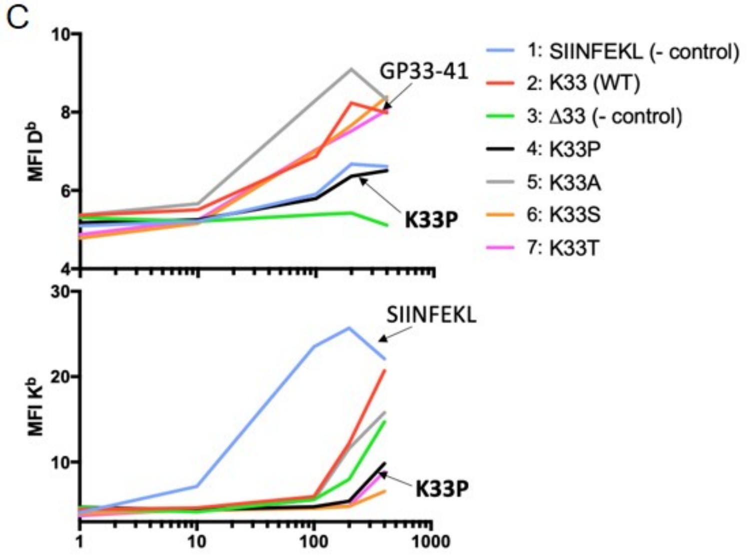


***Figure S4 - Peptides from splice junction in neoantigen module are not presented as antigens***

**a**, The antisense DNA sequence of exon 2 encodes GP34-41 with a preceding amino acid encoded by nCA, which could encode an Ala, Ser, Thr, or Pro residue **b**, the predicted binding of each peptide (SIINFEKL control, GP34-41, or GP33-41 with K33A, K33S, K33T, or K33P mutations). K33P did not bind to H2-$D^b$ or stimulate T cell activation.

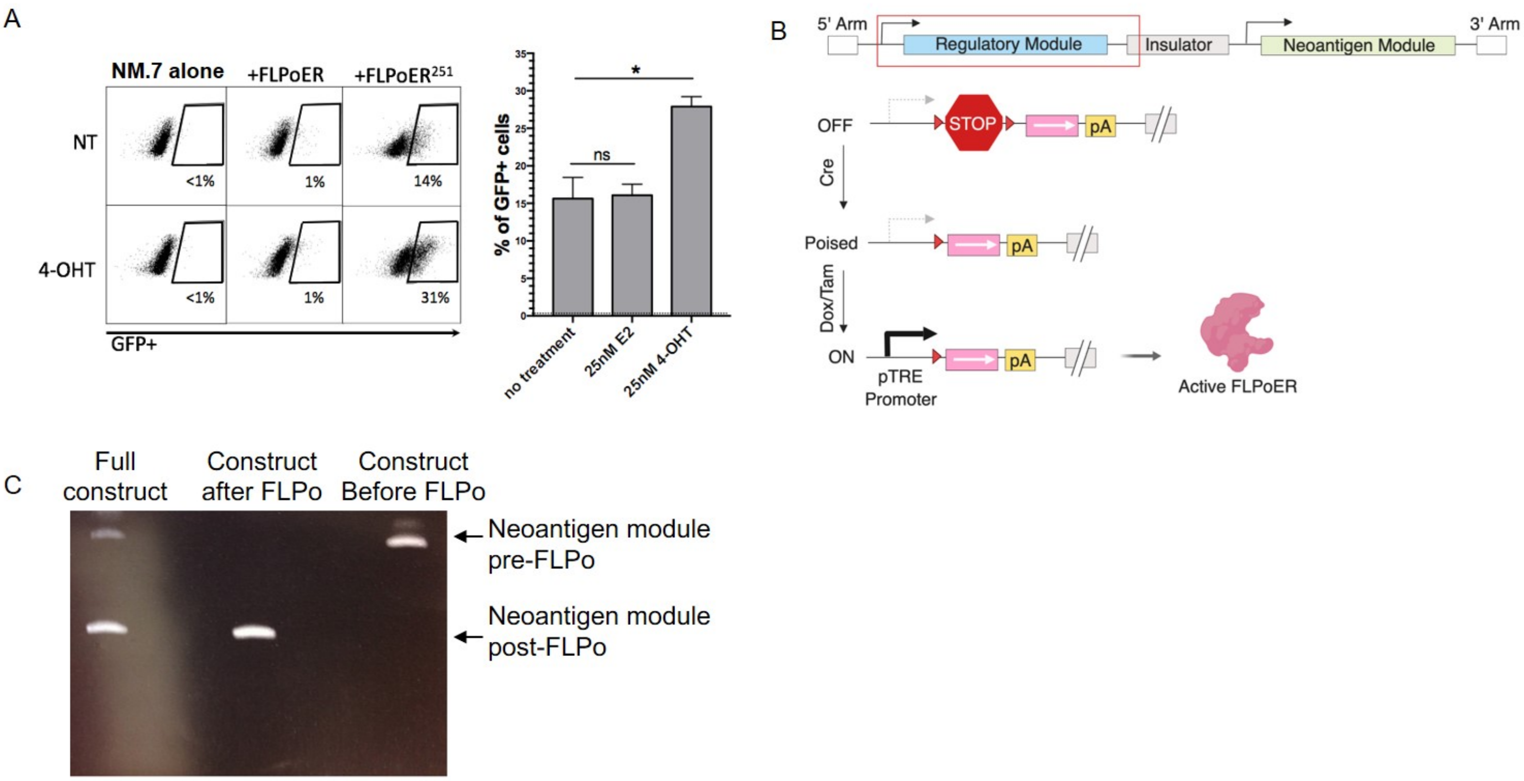


***Figure S5 - Design and development of regulatory module***

**a**, Flow cytometry plots of GFP expression in transiently transfected 293s with either NM.7 alone or in combination with FLPoER or FLPoER$^{251}$. FLPoER$^{251}$, while leakier, is more responsive to 4-OHT treatment than FLPoER and its activity was not increased by estrogen (E2) treatment. **b**, An early regulatory module design, with pTRE:Lox-STOP-Lox (LSL):FLPoER$^{251}$ 2x CGG insulator, and the neoantigen module. We discovered in **c**, that the neoantigen module in this construct was recombined by FLPo activity in *E coli*, which led to the later inverted design.

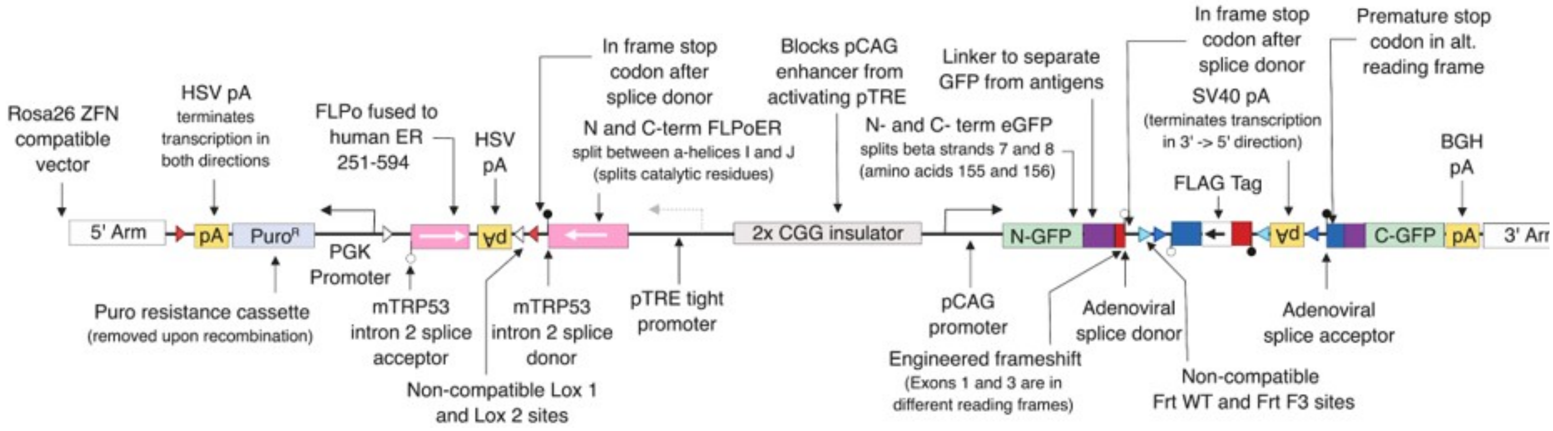


***Figure S6 - Schematic diagram of completed NINJA construct***

Final version of the NINJA construct, including the final regulatory and neoantigen modules.

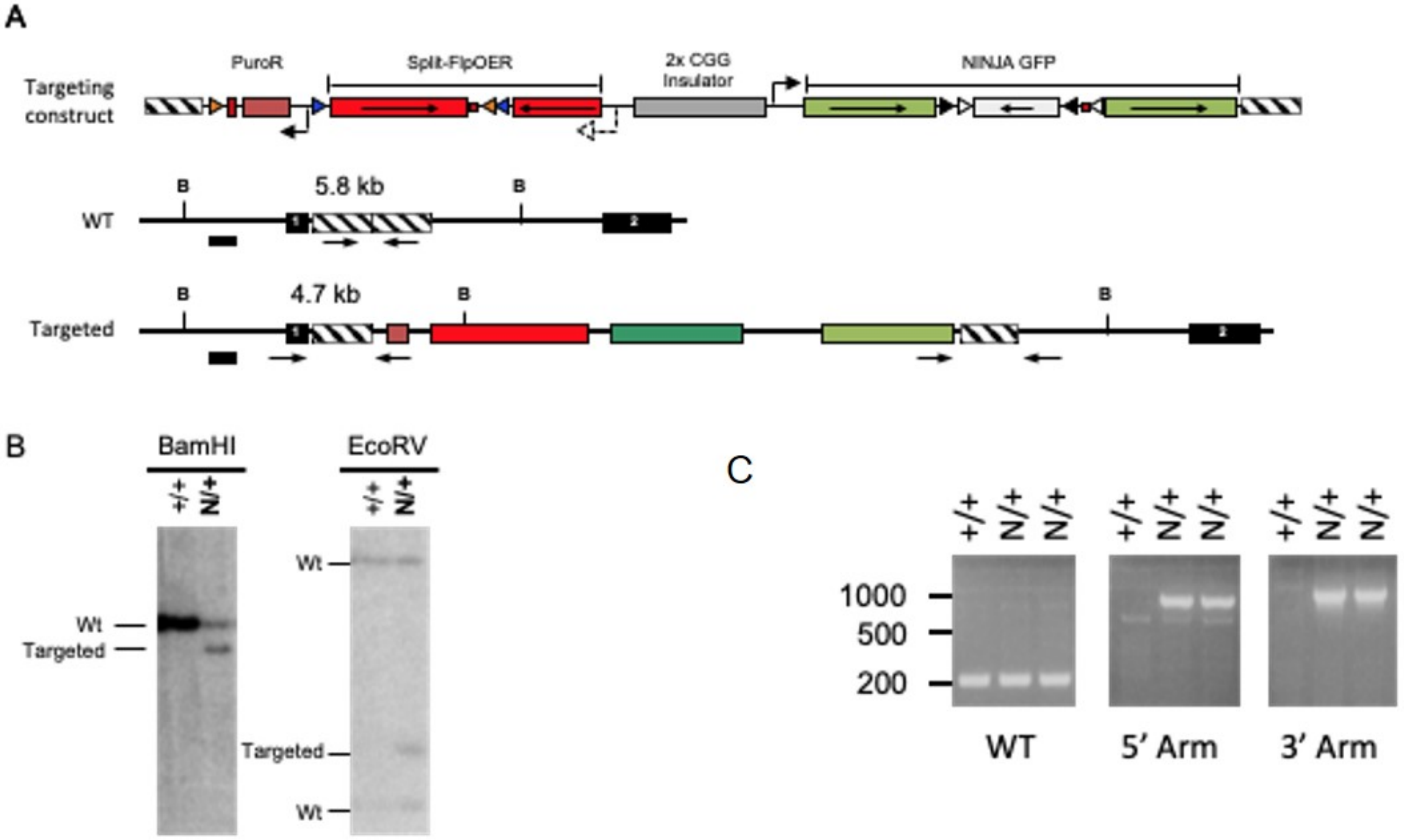


***Figure S7 - Targeting of ES cell clones for NINJA***

**a**, Schematic showing the Rosa 26 targeting construct used for generation of the NINJA mouse. **b**, Southern blotting confirmation of successful target insertion of NINJA **c**, Confirmation of germline transmission in two pups via PCR in the Rosa locus.

A

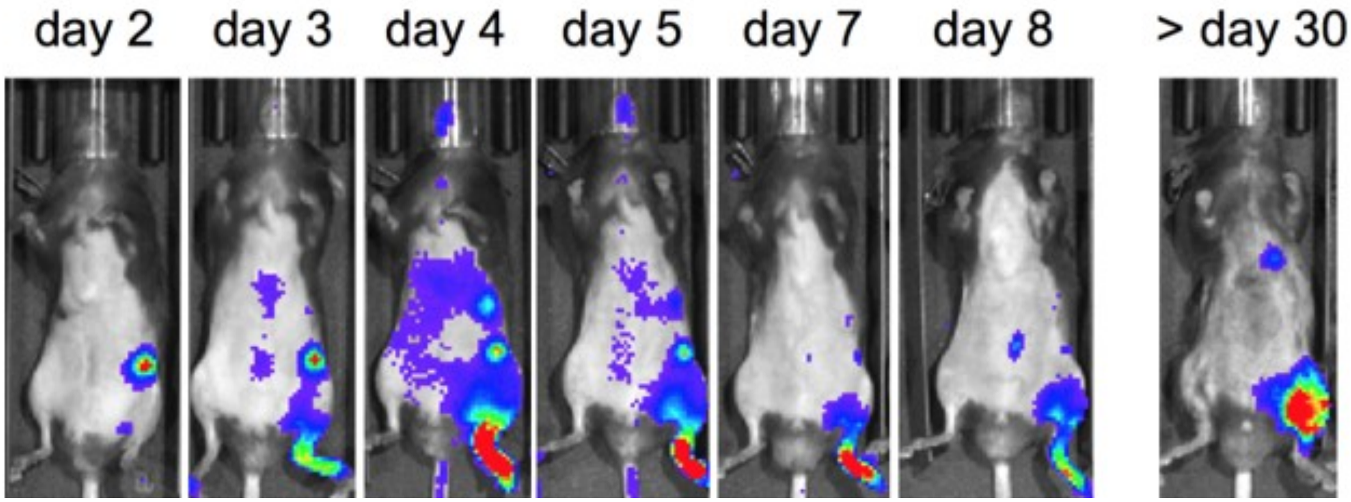


***Figure S8 - Time course of neoantigen-specific CD8 T cell response in NINJA***

Images show local accumulation and expansion of $fLuc^{+}$ P14 cells adoptively transferred into NINJA mice infected S.C. in the footpad with Ad-FLPo ($10^{7}$ PFU/mouse) and imaged by IVIS at the indicated day after infection. Representative mice are shown ($n = 3$). Intensity of signal (blue to red) indicates accumulation of T cells.

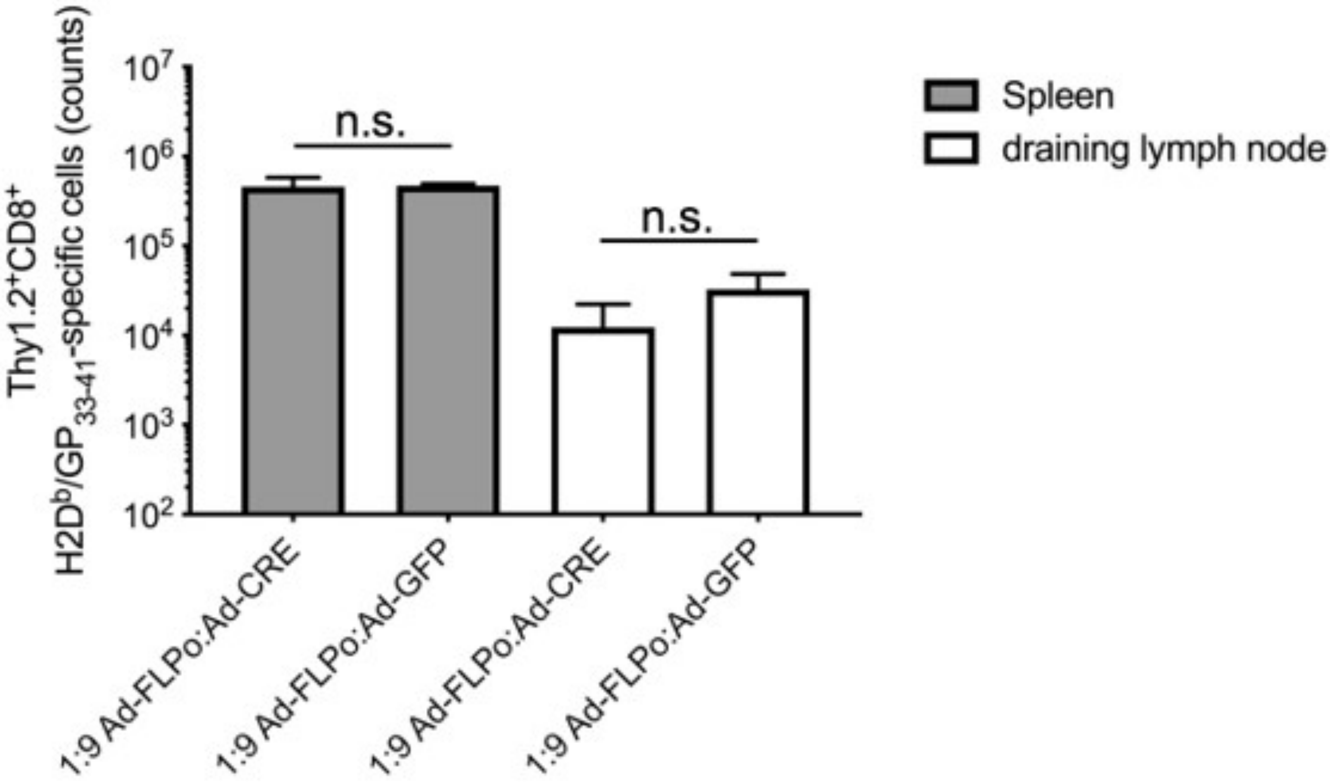


***Figure S9 - Ad-Cre infection does not lead to leaky neoantigen expression***

Quantification of endogenous H2Db/$GP_{33-41}$-specific Thy1.2$^{+}$CD8$^{+}$ cells from the spleen (gray bars) and draining lymph node (white bars) of NINJA mice 8 days after S.C. infection in the footpad with the indicated mixes of Ad-FLPo + Ad-Cre or Ad-FLPo + Ad-GFP (total dose $10^7$ PFU/mouse) was performed by flow cytometry. Representative experiment is shown ($n$ = 3). n.s. = difference not significant by unpaired $t$ test for comparisons of (Ad-FLPo + Ad-Cre) vs. (Ad-FLPo + Ad-GFP). Average values ± SEM are shown.
